# Global patterns in gene content of soil microbiomes emerge from microbial interactions

**DOI:** 10.1101/2023.05.31.542950

**Authors:** Kyle Crocker, Kiseok Keith Lee, Milena Chakraverti-Wuerthwein, Zeqian Li, Mikhail Tikhonov, Madhav Mani, Karna Gowda, Seppe Kuehn

**Affiliations:** Department of Ecology and Evolution, The University of Chicago, Chicago, IL 60637, USA; Center for the Physics of Evolving Systems, The University of Chicago, Chicago, IL 60637, USA; Biophysical Sciences Graduate Program, The University of Chicago, Chicago, IL 60637, USA; Department of Physics, The University of Illinois at Urbana-Champaign, Urbana, IL 61801, USA; Department of Physics, Washington University in St. Louis, St. Louis, MO 63130, USA; Department of Engineering Sciences and Applied Mathematics, Northwestern University, Evanston, IL 60208, USA; Department of Molecular Biosciences, Northwestern University, Evanston, IL 60208, USA; NSF-Simons Center for Quantitative Biology, Northwestern University, Evanston, IL 60208, USA

## Abstract

Microbial metabolism sustains life on Earth. Sequencing surveys of communities in hosts, oceans, and soils have revealed ubiquitous patterns linking the microbes present, the genes they possess, and local environmental conditions. One prominent explanation for these patterns is environmental filtering: local conditions select strains with particular traits. However, filtering assumes ecological interactions do not influence patterns, despite the fact that interactions can and do play an important role in structuring communities. Here, we demonstrate the insufficiency of the environmental filtering hypothesis for explaining global patterns in topsoil microbiomes. Using denitrification as a model system, we find that the abundances of two characteristic genotypes trade-off with pH; *nar* gene abundances increase while *nap* abundances decrease with declining pH. Contradicting the filtering hypothesis, we show that strains possessing the Nar genotype are enriched in low pH conditions but fail to grow alone. Instead, the dominance of Nar genotypes at low pH arises from an ecological interaction with Nap genotypes that alleviates nitrite toxicity. Our study provides a roadmap for dissecting how global associations between environmental variables and gene abundances arise from environmentally modulated community interactions.

## Introduction

The metabolic activity of microbiomes in natural environments from soils to oceans drives the global cycling of carbon, nitrogen, and other elements essential to life on Earth (***?***). This activity arises from the collective action of metabolic pathways carried by diverse, interacting microbes in communities. Despite this complexity, global sequencing surveys of natural communities have revealed ubiquitous patterns relating the abundance of the genes that make up these pathways to local environmental variables. For example, bacterial metabolic capabilities vary with nutrient levels in marine systems (*1*), and gene content changes with soil pH (*2–4*) and host diet (*5–7*). Therefore, variation in the local environment modulates microbiome gene content and metabolic activity upon which the biosphere relies (*3, 8, 9*). Understanding the origins of environmentally-mediated variation in microbiome gene content is a necessity to understand how human activity impacts global nutrient cycles.

How do environmentally dependent patterns in gene content reproducibly emerge in microbial communities across the planet? One common hypothesis is environmental filtering: strains exhibiting traits that are adapted to their local environment are able to colonize that environment even in the absence of any other strains being present (*10*). In this case, the dominance of a strain in an environmental condition is not dependent on biotic interactions, and depends only on the traits of the environmentally selected strains. In some cases, associations between environmental variables and certain genotypes make intuitive sense with the environmental filtering hypothesis. For example, the relative abundance of acid-tolerant Acidobacteria in soil increases as the pH declines (*9,11,12*). Similarly, the prevalence of anaerobic nitrate respiration and anoxygenic photoautotrophy increase with decreasing concentrations of oxygen in the global ocean microbiome (*1*). In these cases, we presume that specific metabolic strategies are enriched in niches where those traits facilitate a competitive advantage. However, we also know that microbial communities are complex systems with strong interactions between constituents (*13–16*), resulting in feedback, counter-intuitive inhibitory effects, evolutionary consequences, and predation, all of which conspire to determine abundances (*17–21*). Thus environmental factors can modulate interactions and drive impacts on community diversity, gene content, and metabolic activity. Despite this, it remains unclear what role these interactions play in giving rise to ubiquitous large-scale patterns observed in communities.

From this vantage, a problem arises: how can simple patterns in environmentally mediated gene content emerge given the apparent complexity of community interactions? One possibility consistent with the environmental filtering hypothesis is that interactions are weak, and changes in gene content reflect an adaptation of individual genotypes to local environmental conditions. A second possibility is that the patterns in gene content emerge from ecological interactions whose strength and specificity are modulated by local environmental variables. In this case, the emergence of reproducible patterns requires that the interactions exhibit regularity across different locations with similar environmental conditions. However, given the complexity of interactions in microbiomes, imagining how this regularity could manifest is a challenge.

Thus, two points are clear. First, patterns relating environmental variation to community composition are ubiquitous in many microbiomes. Second, it is clear from laboratory studies that interactions are important in structuring microbial communities and these interactions can depend on the environmental context. Less direct evidence also strongly suggests that interactions are important in more complex communities in the wild (*22*). What remains unclear is how interactions in very complex communities can give rise to the ubiquitous regular patterns we observe in natural consortia on a large scale. Here, we develop and demonstrate a new approach to answering this question.

Here, we present evidence that global patterns in the gene content of microbial communities emerge from environmentally dependent interactions between members of the microbiome. We use bacterial denitrification as a model process to understand the mechanisms underlying global patterns in gene content. We quantify patterns in the abundances of genes associated with denitrification in the global topsoil microbiome (*3*), finding that soil pH strongly associates with variation in denitrification reductase gene content in soils. Using enrichments and quantitative phenotyping of isolates, we demonstrate the insufficiency of the environmental filtering hypothesis to explain these global patterns. Specifically, we show that strains possessing the denitrification gene Nar (Nar^+^), encoding a cytoplasmic nitrate reductase, are globally enriched in low pH soils, and that strains possessing this gene dominate low pH enrichment cultures as well. Counterintuitively, Nar^+^ strains generically accumulate nitrite, which is toxic in acidic conditions, giving rise to reduced growth when cultured alone. We show that a strong interaction between Nar^+^ strains and strains possessing the alternative nitrate reductase Nap reduces this toxicity, allowing Nar^+^ to dominate at low pH. We argue that these interactions yield reproducible genomic patterns on a global scale as a result of the conserved physiological properties of the genotypes involved (*23*) and the interactions these physiological traits support. We support this claim by showing that denitrifying populations in soils are dominated by strains that are taxonomically similar to the enriched isolates used in our study. Further, a soil microcosm experiment reveals that just a handful of taxa out of the thousands present in soils respond to the presence of nitrate, suggesting that interactions between a small number of strains may be relevant even in the ecological context of a soil microbiome. Our study provides a unified way of understanding ubiquitous patterns in microbiomes as emerging from environmentally-modulated interactions between members of the community.

## Results

### Quantifying global patterns in topsoil microbiome gene content

We first sought to quantify variation in metabolic gene content on a global scale. To accomplish this, we focused on the soil microbiome because of the importance of these environments for global biogeochemistry, and the fact that global metagenomic surveys of topsoil microbiomes have recently been completed (*3*). Within the soil microbiome, we studied the process of denitrification, because it is ubiquitous, it arises from enzymes that are well-studied and reliably annotated (*24, 25*), and is performed by readily culturable taxa (*23, 26, 27*). As we demonstrate below, these properties enabled us to reliably identify genomic patterns in gene content from sequencing data and dissect interactions using wild isolates.

In soils, denitrifiers are present at substantial abundances (∼1 × 10^6^ cells/g, Ref. 28). These bacteria drive a cascade of reduction reactions (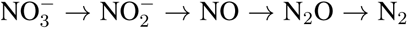, Fig. 1A) to utilize organic carbon anaerobically via respiration. The cascade of reduction reactions is carried out by phylogenetically diverse bacteria (*29*), utilizing specific reductases to perform each reaction (Fig. 1A). The structure of the denitrification pathway varies between taxa, with some strains performing all four steps of the cascade, and others performing nearly arbitrary subsets of the four reactions (*27*). As such, denitrification is often a collective process in the wild.

**Figure 1:**
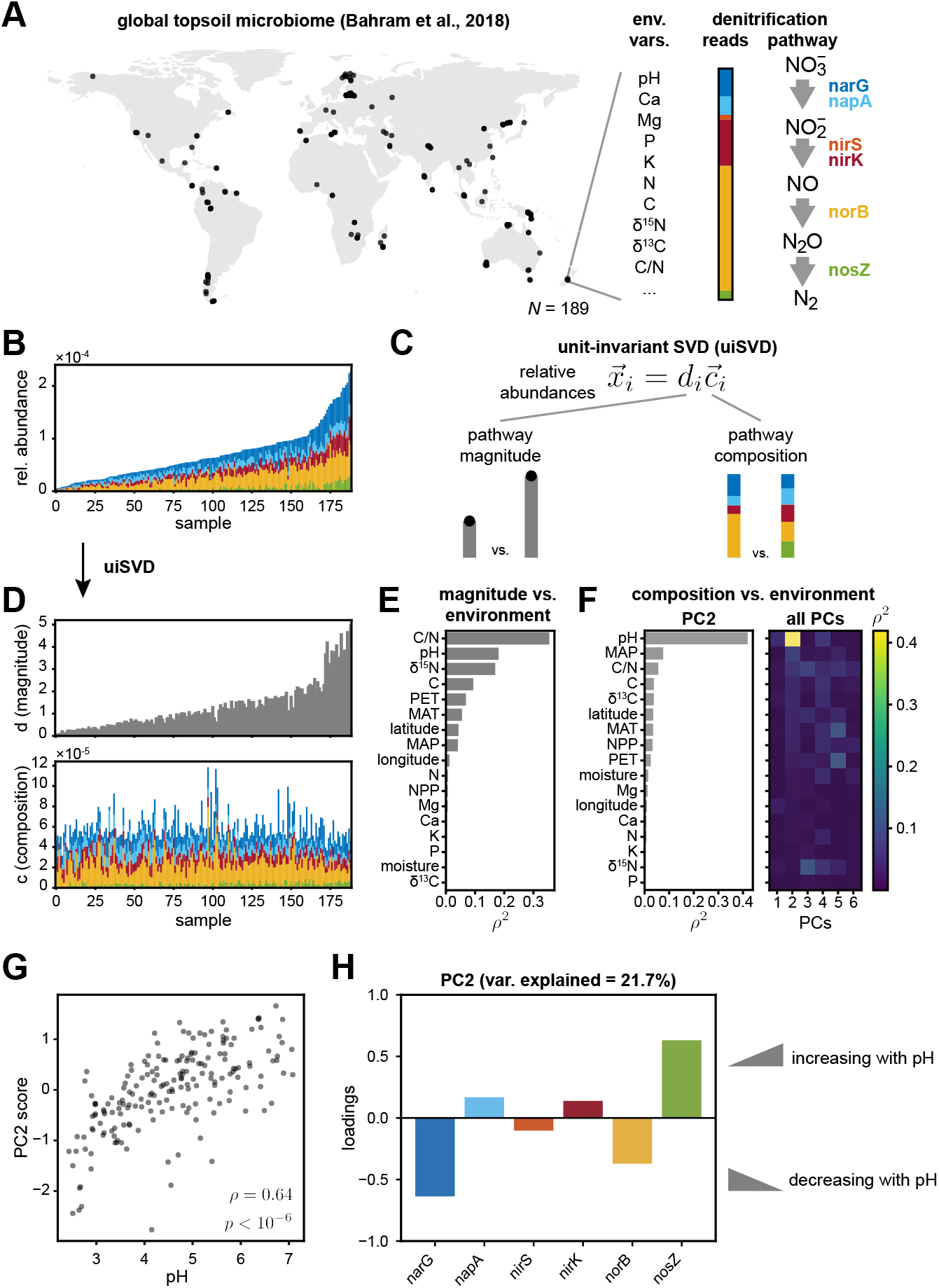
pH is associated with covariation in denitrification pathway composition in the global topsoil microbiome. **(A)** Topsoils sampled at *n* = 189 globally-distributed sites were chemically characterized (pH, Ca, Mg, …), sequenced via shotgun metagenomics, and functionally annotated by Bahram et al. (*3*). **(B)** The relative abundance of denitrification reductases in each soil sample (relative to total gene content) are plotted in order of increasing total relative abundance. Reductase color legend indicated in panel A. **(C-D)** Unit-invariant singular value decomposition (uiSVD) (*30*) was used to decompose the data in panel B into contributions due to pathway magnitude (*d_i_*) and pathway composition (*c̃_i_*). The results of this decomposition are plotted in panel **D**. **(E-F)** Pathway magnitude values and principal component (PC) scores for pathway composition obtained via uiSVD (Methods) are compared with 17 environmental variables, and squared Pearson correlation coefficients are shown. Pathway magnitude is most correlated with C/N ratio (*ρ*= *-*0.59, *p<* 10*^-^*^6^ via one-tailed randomization test; Fig. S1), while scores of PC2 are most correlated with pH (*ρ*= 0.64, *p <* 10*^-^*^6^ via one-tailed randomization test; **G**). **(H)** Loadings of PC2 are shown, where positive values indicate reductase content that increases with pH, and vice versa. See also Figs. S1, S2.

To study denitrification gene content in soils we utilized the Bahram et al. (*3*) survey of the topsoil microbiomes. The dataset comprises *n* = 189 samples taken from sites across the planet (Fig. 1A; Ref. 3). Crucially, this dataset combines shotgun metagenomics with a detailed characterization of the physicochemical properties of each soil sample, including pH, ion concentrations, nutrient levels, and site climate characteristics (Methods). The fact that these measurements were performed in a standardized fashion on soils from across the globe enabled us to relate environmental factors to the abundances of genes encoding denitrification enzymes.

First, we quantified variation in denitrification gene content across soils in the global topsoil microbiome. To do this, we computed the fraction of reads in each sample that map to any of the six main reductases in the denitrification pathway (*narG*, *napA*, *nirS*, *nirK*, *norB*, and *nosZ*; Fig. 1A). Note, due to the relatively low sequencing depth of these samples, we were not able to construct reliable metagenome-assembled genomes, so the broader genomic context of these genes is not known. The result is a table that contains the fraction of all reads mapping to any of the reductases in the denitrification pathway (vertical axis, Fig. 1B). We refer to this total fraction as the “magnitude” of the pathway (Fig. 1C). Our analysis also quantifies the pathway composition, defined as the relative contribution of each reductase to the total (colors, Fig. 1B). Denitrification pathway magnitude varies by about 50-fold across soil samples, while pathway composition also varies from sample to sample, but this variation is challenging to visually distinguish from variation in pathway magnitude (Fig. 1B).

To disentangle pathway magnitude from composition, we developed a new approach based on unit-invariant singular value decomposition (uiSVD, (*30*); Methods). uiSVD decomposes the relative abundances of denitrification genes in each sample (*x⃗_i_*) into a contribution arising from the magnitude of the pathway (*d_i_*) and the pathway composition (*c̃_i_*) (Fig. 1C). The result of applying this method to the denitrification reductase relative abundance matrix is shown in Fig. 1D. Note that the data were not logratio-transformed to account for compositionality, as the high diversity of the gene abundance dataset makes the effects of compositionality negligible (Methods). Next, we sought to determine how environmental variables are associated with the magnitude and composition of the denitrification pathway in soils.

### pH correlates with pathway composition on a global scale

To connect environmental variation to patterns in pathway magnitude, we examined correlations between environmental variables and magnitude. We considered squared Pearson correlations (*ρ*^2^), across the *n* = 189 soil samples, between pathway magnitudes (*d_i_*) and each environmental variable in turn. We found that the magnitude of the pathway was most strongly associated with the C/N ratio (*ρ*^2^ = 0.35 Fig. 1E). The correlation is negative (*ρ* = *-*0.59, *p <* 10*^-^*^6^ via one-tailed randomization test; Fig. S1), consistent with previous work showing that denitrification is enhanced at low C/N ratios (*31*).

To understand how environmental variation impacts the prevalence of the different denitrification enzymes, we computed the principal components (PCs) of the composition matrix (Methods, Fig. S2), which represent covarying groups of genes. By looking at correlations between PC scores (projections of composition onto the PCs) to the environmental variables, we found that a single component of variation, PC2 (21.7% variance explained), was associated with pH much more strongly than any other combination of components and environmental variables (*ρ* = 0.64, *p<* 10*^-^*^6^ via one-tailed randomization test; Fig. 1F, G). Examining the loadings of each reductase in PC2 (Fig. 1H) reveals how denitrification reductase genes vary with pH on a global scale: some reductase genes are enriched as pH decreases (*narG*, *nirS*, *norB*), while others decline as pH decreases (*napA*, *nirK*, *nosZ*). While pH is widely appreciated as a strong contributor to community diversity in soils (*8*), and pH is known to be important for denitrification (*32, 33*), our analysis provides a quantitative view of how pH impacts which genes are involved in denitrification on a global scale. Our approach provides a principled, generalizable route to dissecting how environmental variation drives compositional changes in microbiomes.

What causes some gene abundances to rise and others to fall with changes in pH? A strict environmental filtering hypothesis would suggest that strains possessing specific reductases are favored in soils of different pH. Under this assumption, Fig. 1F advances the hypothesis that strains possessing the nitrate reductase Nar are favored at low pH, while those possessing the nitrate reductase Nap are favored at neutral pH. However, to our knowledge there is no direct evidence for this hypothesis, nor is this hypothesis easily tested from sequencing data alone. Alternatively, it is also possible that pH is not causally associated with variation in reductase content, but rather related to unmeasured causally-associated variables. Therefore, we sought to test the dependence of denitrification genotypes on pH experimentally using an enrichment culture approach.

### Enrichment cultures recapitulate global patterns in denitrification gene content

We reasoned that if pH is causally related to genotype abundances as shown in Fig. 1F, we should be able to recapitulate these patterns in the lab. To experimentally test the associations between pH and denitrification genotypes, we performed enrichment cultures under different pH conditions (Fig. 2A). We extracted diverse microbial communities from six soil samples (Table S1), and inoculated them into chemically defined media buffered at pH 6.0 and 7.3 (growth slows significantly below pH 6.0). We then passaged these communities serially for 12 total 72 h growth cycles, supplying 1.75 mM nitrate at the beginning of each cycle. We chose these conditions because pH in soils is often strongly buffered (*34–36*), mM concentrations of nitrate are often present (*37, 38*), and the growth medium was developed to capture a diverse range of denitrifiers (*26*). The resulting enrichment cultures were then sequenced to assay taxonomic and functional metagenomic composition. The details of all culturing and sequencing procedures can be found in Methods.

**Figure 2:**
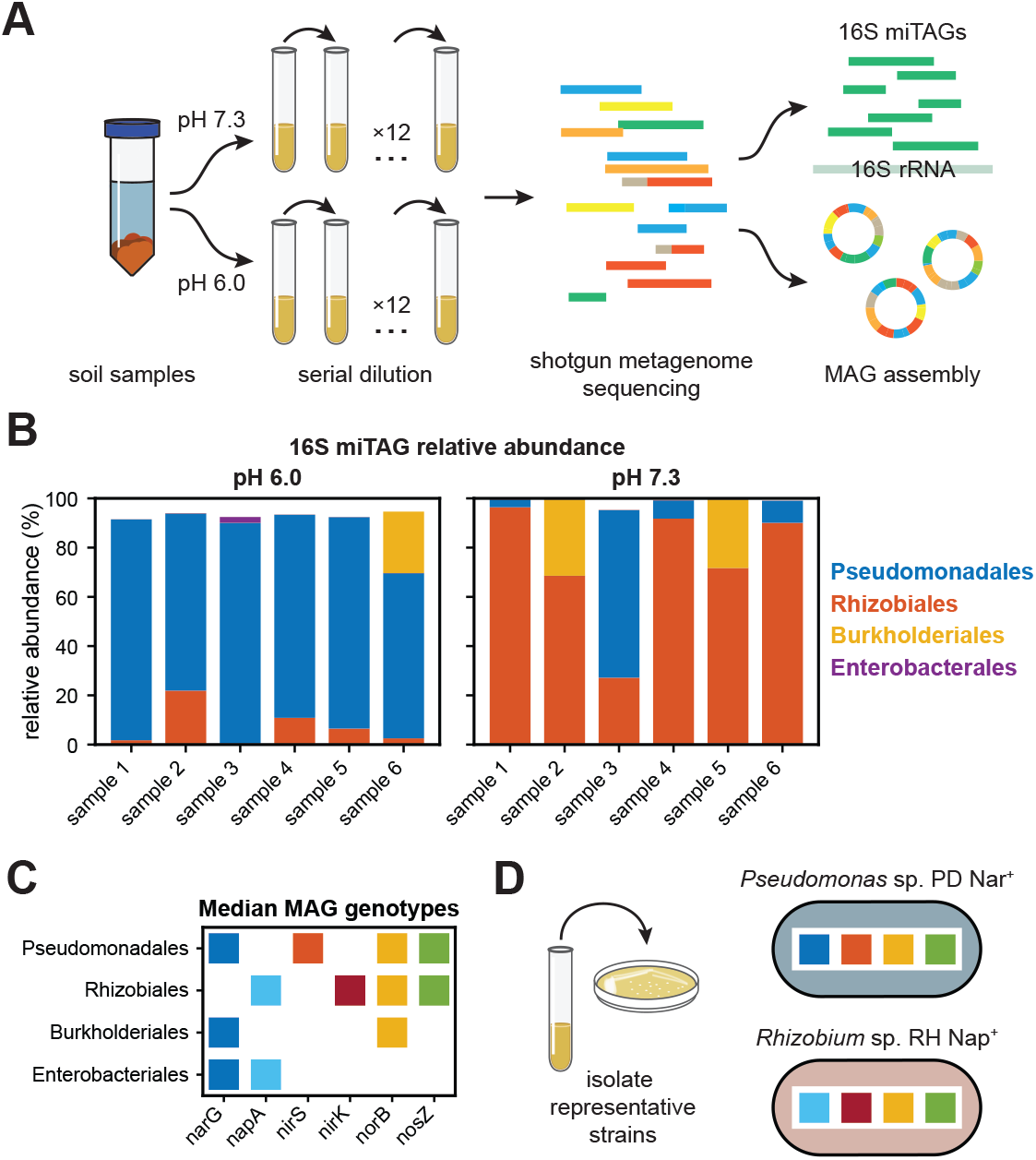
Enrichment cultures recapitulate global patterns in denitrification gene content. **(A)** Denitrifying communities were enriched from soil samples in two pH conditions (6.0 and 7.3), and then sequenced via shotgun metagenomics to measure taxonomic composition and genotypes. Six soil samples were mechanically homogenized and used to inoculate a serial dilution experiment in denitrifying (anaerobic) conditions. After 72 h growth cycles, cultures were repeatedly passaged (12*⇥*) into a defined medium containing 2 mM nitrate via a 1/8 dilution factor (Methods). **(B)** Endpoint community compositions are shown at the level of taxonomic order for enrichments in pH 6.0 and 7.3. Compositions were determined by extracting 16S rRNA fragments (miTAGs) from shotgun metagenome data (*39*), and then taxonomically annotating miTAGs via RDP (*40*). Taxa present at a relative abundance less than 1% are omitted. **(C)** Median denitrification reductase genotypes are shown for metagenome-assembled genomes (MAGs) corresponding to the four most abundant taxa present in the composition data in panel B. MAGs were functionally annotated using RAST (*41*), and the median genotype was computed over MAGs obtained in different samples. **(D)** Strains were isolated from cryopreserved samples from the enrichment endpoint (Methods). The strain *Pseudomonas* sp. PD Nar^+^ represents the dominant taxa present across samples at pH 6.0, and *Rhizobiales* sp. RH Nap^+^ represents the dominant taxa across samples at pH 7.3. See also Figs. S3, S4.

First, to assay taxonomic composition of the endpoint enrichment cultures, we extracted 16S rRNA miTAGs (*39*) from shotgun metagenome sequencing data and annotated them (*42*) (Methods). This analysis revealed that endpoint enrichments were dominated by taxa from the orders Pseudomonadales and Rhizobiales, with the former comprising the majority of all six enrichments at pH 6.0 and the latter making up the majority of five out of six enrichments at pH 7.3 (Fig. 2B). Because the starting points of enrichments at both pH conditions were the same, this result suggests that pH 6.0 selects for Pseudomonadales strains, while pH 7.3 selects for Rhizobiales strains. Taxa from the orders Burkholderiales and Enterobacteriales were present at comparably lower levels, and no other taxonomic orders were present in any enrichment at an abundance greater than 1%.

To determine the genotypes of taxa in the enrichment experiments, we constructed metagenome-assembled genomes (MAGs) (*43*) (Methods). Annotating these MAGs revealed that taxa from different enrichments had largely similar denitrification genotypes (Fig. 2C); in particular, Pseudomonadales MAGs typically encoded the reductases NarG and NirS, while Rhizobiales MAGs encoded the NapA and NirK reductases. A second enrichment experiment, performed with seven additional soil samples enriched at a broader range of pH values, also recapitulated the same basic pattern in genotype abundances, with taxa possessing NarG heavily enriched at pH 5.0 and 5.5 (Fig. S4; Methods).

Strikingly, these genotypes recapitulate the patterns in denitrification gene content observed in the topsoil microbiome (Fig. 1F); namely, our enrichment resulted in a greater prevalence of *narG* and *nirS* at pH 6.0 (blue bars in Fig. 2B, corresponding to the top row of Fig. 2C), and alternatively a greater prevalence of *napA* and *nirK* at pH 7.3 (orange bars in Fig. 2B, corresponding to the second row of Fig. 2C). This agreement between the enrichment and our analysis of the global topsoil microbiome suggests that (1) pH is likely causally related to the patterns of gene covariation described in Fig. 1F, and (2) the mechanisms shaping these patterns in nature are also at play *in vitro*. Altogether, this points to the possibility that we can uncover the mechanistic basis for global patterns of gene-environment covariation by interrogating our enrichment experiments of community dynamics and assembly.

### Environmental filtering does not explain outcome of acidic enrichments

To uncover the mechanisms that give rise to the observed pattern between denitrification gene content and soil pH (Fig. 1), we isolated taxa from the endpoint enrichment experiment from the genera *Pseudomonas* and *Rhizobium* (Fig. 2D, Methods). These strains correspond to the Nar^+^ and Nap^+^ genotypes, respectively (Fig. 2D). We denote the taxonomy and genotype of these isolates as PD Nar^+^ and RH Nap^+^.

The environmental filtering hypothesis would suggest that the Nar^+^ genotype should thrive in acidic conditions and the Nap^+^ genotype in neutral conditions. To test this hypothesis, we performed a serial dilution experiment with strains grown in isolation (Methods). We measured the temporal dynamics of nitrate, nitrite, and total biomass at the end of each cycle in this experiment (Fig. 3).

**Figure 3:**
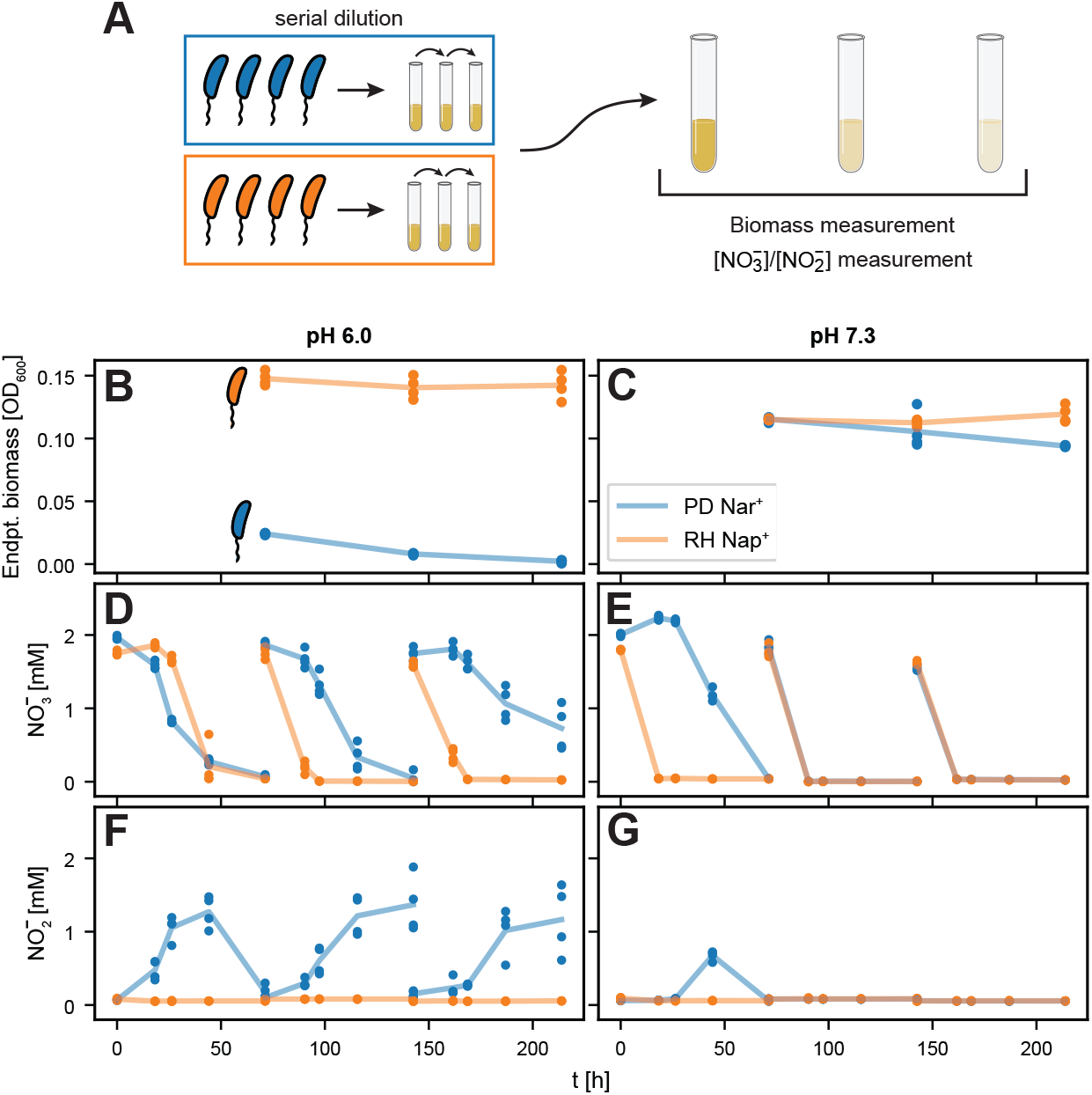
Environmental filtering does not explain outcome of acidic enrichments. **(A)** Schematic of the experimental design. PD Nar^+^ (blue) and RH Nap^+^ (orange) were passaged repeatedly in monoculture under denitrifying (anaerobic) conditions. After each 72 h growth cycle, cultures were passaged (3x) into a defined medium containing 2 mM nitrate using a 1*/*8 dilution factor. Biomass is measured via 600 nm absorbance at the end of each cycle, and nitrate and nitrite concentrations are measured throughout each cycle using a Griess assay (Methods). (**B** and **C**) Endpoint biomass is shown for each strain at both pH 6.0 (panel **B**) and 7.3 (panel **C**). Lines connect the average across replicates. RH Nap^+^ (orange) produces more biomass at both pH levels, while PD Nar^+^ (blue) appears to decay in abundance at pH 6.0 (despite PD Nar^+^ community enrichment in this condition, Fig. 2B). (**D-G**) Nitrate and nitrite concentration dynamics are shown at six timepoints throughout each cycle in each condition. PD Nar^+^ nitrate and nitrite reduction rates slow at pH 6.0 as the growth-dilution cycles progress, consistent with its reduction in biomass (Panels **D** & **F**, blue). PD Nar^+^ (blue) accumulates nitrite at each pH condition, whereas RH Nap^+^ (orange) does not (Panels **F** & **G**). Technical replicates (*n* = 4) are shown for each strain in each experimental condition, with lines connecting the averages of these replicates.

In line with the environmental filtering hypothesis, the strain RH Nap^+^ (representative of taxa most abundant in pH 7.3 enrichments) was more fit than PD Nar^+^ at pH 7.3. In particular, RH Nap^+^ produced more biomass in the final cycle at pH 7.3 (at t = 213 h: RH Nap^+^ OD_600_ = 0.119*±*0.006, while PD Nar^+^ OD_600_ = 0.094 *±* 0.001; Fig. 3C) and accumulated less nitrite in the first cycle (at t = 44 h: 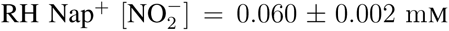, while 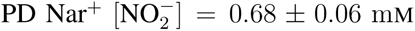; Fig. 3G), despite reducing nitrate more quickly (for t *<* 72 h: orange data values are less than blue data values in Fig. 3E). Additionally, simulated competition between these strains confirms that RH Nap^+^ is expected to be enriched at pH 7.3 based on monoculture phenotype. Briefly, a consumer-resource model was parameterized following (*44*) and used to simulate coculture growth of the two strains across batch culture cycles. This simulation predicted that RH Nap^+^ relative abundance would approach 1 on the timescale of the enrichment experiments (Supplementary Information).

In contrast, results at pH 6.0 were not consistent with the environmental filtering hypothesis. PD Nar^+^ produced only a small amount of biomass, with levels decreasing with each cycle, suggestive of cell death (Fig. 3B). At the same time, this strain reduced nitrate more slowly than RH Nap^+^ in cycles 2–4 and reduced nitrite more slowly in all cycles (Fig. 3D, F). Crucially, PD Nar^+^ transiently accumulated nitrite (blue curves, Fig. 3F), with accumulation increasing over successive cycles (Fig. 3F).

These experiments reveal a paradox at pH 6.0: PD Nar^+^ dominated in a community context (Fig. 2B) and yet performed poorly by itself. This result is inconsistent with the environmental filtering hypothesis, suggesting that the dominance of Nar^+^ genotypes in acidic enrichment experiments emerged from community-level processes.

### Nitrite toxicity impacts denitrification activity of isolates at low pH

As a route to understanding how community-level processes account for the discrepancy between soil enrichments (Fig. 2) and monocultures (Fig. 3), we next sought to uncover the mechanisms underlying the poor performance of PD Nar^+^ monocultures at pH 6.0. Given the accumulation of nitrite in this experiment (Fig. 3F), as well as literature indicating the toxicity of nitrite under acidic conditions (*45–47*), we hypothesized that nitrite accumulation inhibited the growth of PD Nar^+^ at pH 6.0.

We tested whether nitrite inhibited the metabolic activity of either isolate at pH 6.0. To do this, we initiated monocultures of each strain at pH 6.0, supplying 1.75 mM nitrate and varying concentrations of nitrite. We then measured nitrate and nitrite dynamics over 72 h of growth (Fig. 4A).

**Figure 4:**
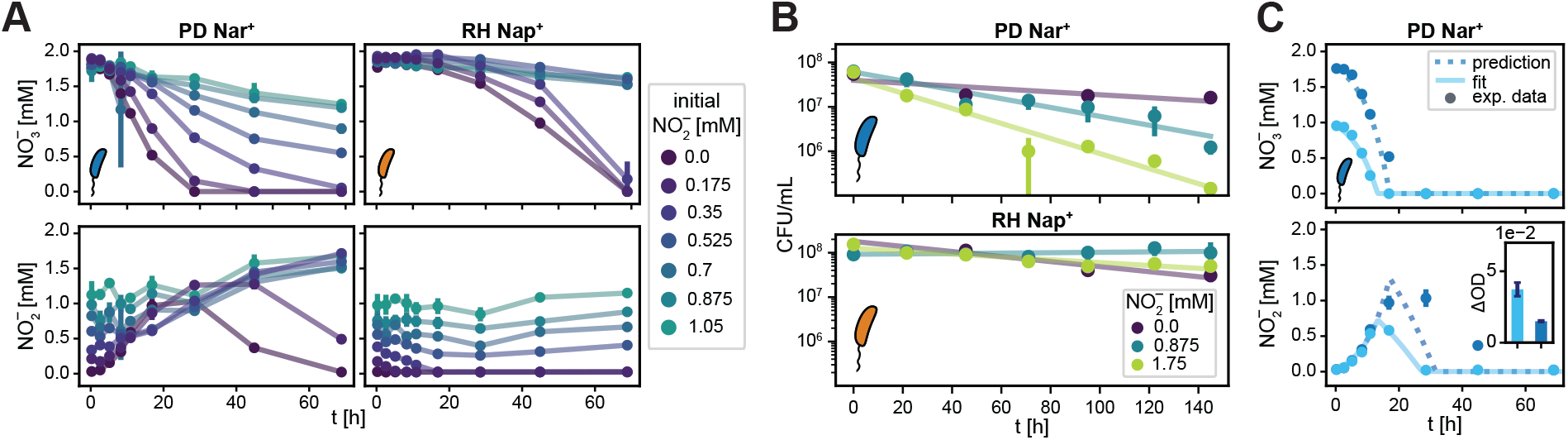
Nitrite toxicity impacts denitrification activity of isolates at low pH. (**A**) PD Nar^+^ and RH Nap^+^ were inoculated in anaerobic monoculture at pH 6.0 with 1.75 mM nitrate and varying nitrite levels (colorbar). Nitrate and nitrite concentration dynamics, shown for each strain, were measured via Griess assay (Methods). As the initial nitrite concentrations 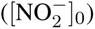 rose, the growth of both strains was inhibited. PD Nar^+^ was unable to fully reduce nitrate when 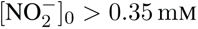 (blue and green curves, top left panel) and was unable to fully reduce nitrite when 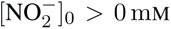 (all except dark purple curve, bottom left panel). Similarly, RH Nap^+^ was unable to fully reduce either nitrate or nitrite when 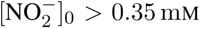 (blue and green curves, right panels). Mean and standard deviation of technical replicates (*n* = 3) are shown. (**B**) PD Nar^+^ and RH Nap^+^ were again grown in anaerobic monoculture at pH 6.0 with varying nitrite levels, but nitrate was not supplied to prevent growth (Methods). Density of colony forming units was measured via plating with replicates (*n* = 3) for each condition, with points indicating means across replicates. Error bars were calculated by weighting across dilution levels, as described in Methods. Lines are log-linear fits, and the (negative) slope of the line indicates mortality rate (Methods). For PD Nar^+^, inferred death rates were 0.008 *±* 0.004, 0.023 *±* 0.004, and 0.039 *±* 0.004 h*^-^*^1^ for 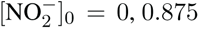, and 1.75 mM, respectively. For RH Nap^+^, inferred death rates were 0.013 *±* 0.004, *-*0.001 *±* 0.001, and 0.007 *±* 0.001 h*^-^*^1^ for 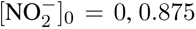, and 1.75 mM, respectively. Uncertainties indicate standard deviations of log-linear fit parameters. (**C**) Nitrate (top) and nitrite (bottom) metabolite dynamics for monocultures of PD Nar^+^, supplied with initial nitrate concentrations of 0.875 mM (light blue) and 1.75 mM (dark blue). A consumer resource model was fit to the low initial nitrate data (solid light blue lines) and used to predict high initial nitrate data (dashed dark blue lines). The prediction failed at intermediate time points (bottom panel), indicating that the accumulation of nitrite slowed the metabolic rate. Inset in bottom panel shows final optical density (OD). Endpoint biomass levels were greater in the 0.875 mM nitrate condition than in the 1.75 mM condition, suggestive of mortality induced by nitrite accumulation. Mean and standard deviation of technical replicates (*n* = 3) of the data are shown.

We found that the extent of inhibition increased with initial nitrite concentration for both strains (Fig. 4A). Interestingly, nitrite appeared to inhibit these two strains in qualitatively different ways. The nitrate reduction rate for PD Nar^+^ slowed in a manner that was continuous with initial nitrite concentration (Fig. 4A), while nitrate reduction rate for RH Nap^+^ showed a threshold-like dependence on initial nitrite concentration 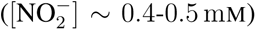. Altogether, these results affirm that the accumulation of nitrite under acidic conditions (blue curves, Fig. 3F) should particularly impact PD Nar^+^, while in contrast, RH Nap^+^ does not accumulate nitrite appreciably (orange curves, Fig. 3F) and therefore should avoid inhibition.

Next, we sought to determine whether nitrite toxicity caused cell death. To do this, we initialized monocultures at pH 6.0, with no nitrate, and 0, 0.875, or 1.75 mM nitrite (Methods). In these conditions, we observed no cell growth, allowing us to quantify mortality via plating (Methods).

The density of colony forming units (CFU/mL) as a function of time is shown for both strains in Fig. 4B. The dark purple line (0 mM nitrite) is a control for stationary-phase mortality in the absence of nitrite. Cell densities below these values indicate death due to nitrite toxicity. We observed the mortality of PD Nar^+^ increased sharply with increasing nitrite concentration at pH 6.0, while the mortality rate of RH Nap^+^ changes little with nitrite concentration. This result strongly suggests that the accumulation of nitrite by PD Nar^+^ in monoculture (Fig. 3F) drives cell death.

Finally, we compared the growth of PD Nar^+^ in pH 6.0 monocultures supplied with different initial concentrations of nitrate. Because nitrite is the direct product of nitrate reduction, lower initial concentrations of nitrate result in less nitrite accumulation, and in principle less toxicity under acidic conditions. As expected, experiments initialized with either 1.75 mM (high) or 0.875 mM (low) nitrate indeed resulted in different degrees of nitrite accumulation (light/dark blue data points, Fig. 4C). Measuring endpoint biomass strongly indicated that the degree of toxicity was greater in the high nitrate condition. PD Nar^+^ produced substantially less biomass in the high nitrate condition (inset, Fig. 4C), amounting to one-fifth of the biomass yield per unit substrate consumed compared with the low nitrate condition.

To further quantify the effect of nitrite toxicity in these experiments, we fit a consumer-resource model (CRM) to the metabolite dynamics and final biomass in the low nitrate condition. The CRM quantifies metabolite dynamics and growth using a Monod equation formalism and has been shown to describe denitrification dynamics reliably (*23*). We fit CRM parameters to the low nitrate condition and used these parameters to predict the high nitrate condition (Methods). If PD Nar^+^ experienced the same degree of toxicity in both conditions, then the fits of the CRM to the low nitrate condition should accurately predict the high nitrate condition. In contrast, a greater degree of toxicity in the high nitrate condition would be reflected in poor model predictions. Indeed, we observed that predictions of the high nitrate condition were poor (dark blue, Fig. 4C); nitrate was consumed more slowly than predicted, indicating a greater degree of toxicity in the high nitrate condition.

Overall, these three experiments demonstrated that nitrite accumulation has a significant deleterious effect on the growth and metabolic activity of PD Nar^+^.

### Community alleviates nitrite toxicity under acidic conditions

PD Nar^+^ failed to grow in monoculture at pH 6.0 (Fig. 3) due to toxicity from accumulating nitrite (Fig. 4), but still managed to dominate the community during enrichment in these conditions (Fig. 2). We hypothesized that since RH Nap^+^ avoids nitrite accumulation in monoculture (dark purple traces, Fig. 4A; orange traces, Fig. 3F), its presence in a co-culture with PD Nar^+^ could serve to reduce nitrite accumulation, alleviating toxicity and allowing PD Nar^+^ to dominate.

To test the hypothesis that the presence of RH Nap^+^ in co-culture alleviates toxicity, we grew a 1:1 mixture (by biomass) of PD Nar^+^ and RH Nap^+^ at pH 6.0, supplying 1.75 mM nitrate (Methods). We found this co-culture accumulated substantially less nitrite relative to the monoculture of PD Nar^+^, producing significantly more biomass (inset, Fig. 5B). Remarkably, nitrite levels in co-culture reached ∼0.5 mM, which is at or below the toxicity threshold identified in earlier experiments (Fig. 4A).

**Figure 5:**
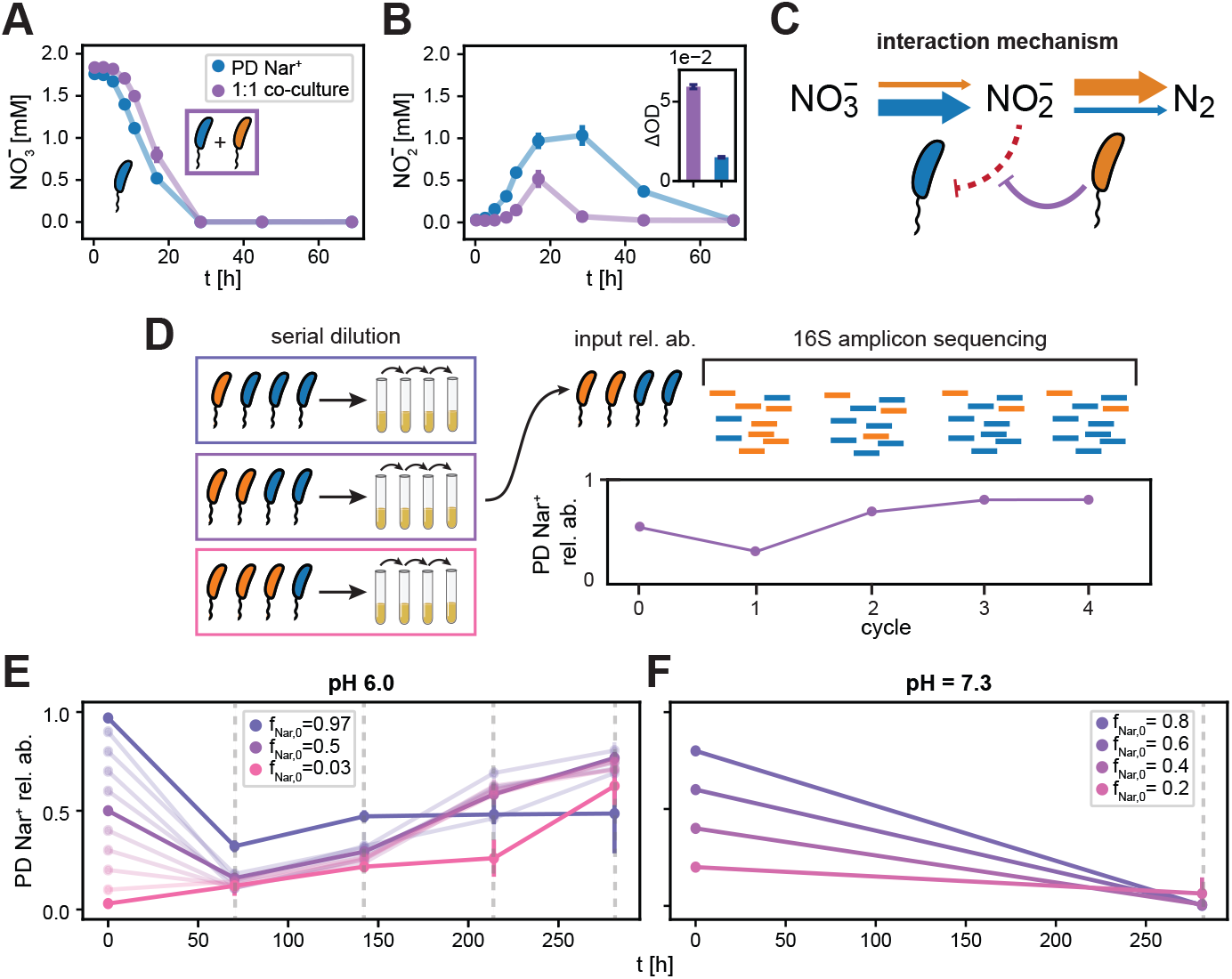
Community alleviates nitrite toxicity under acidic conditions. (**A-B**) Monoculture PD Nar^+^ growth metabolite dynamics (shown previously in Fig. 4C) compared with the those of a 1:1 PD Nar^+^ and RH Nap^+^ co-culture at pH 6.0. Panel **A** shows nitrate concentration and **B** shows nitrite concentration. Initial biomass (OD_600_ = 0.01) and nitrate (1.75 mM) was the same in each condition, but the co-culture (purple) exhibited significantly less nitrite accumulation despite only slightly decreased nitrate reduction rate, as well as significantly higher biomass production (purple bar, inset, panel B). Points and error bars indicate the means and standard deviations, respectively, across technical replicates (*n* = 3). (**C**) Schematic of proposed interaction between PD Nar^+^ and RH Nap^+^. PD Nar^+^ (blue cell) reduces nitrate quickly and accumulates nitrite (thick blue arrow). In monoculture, this nitrite accumulation is self-inhibitory (dashed red line). In co-culture, however, RH Nap^+^ (orange cell) quickly reduces nitrite (thick orange arrow), alleviating the inhibitory effects of nitrite on PD Nar^+^ (purple line). (**D**) Schematic illustrating the multi-cycle co-culture experiment shown in panels **E** and **F**. Mixtures of PD Nar^+^ and RH Nap^+^ were prepared across a range of ratios spanning 0.03:0.97 to 0.97:0.03 (distinguished by color), with total biomass held constant. These mixtures were then transferred to fresh media buffered at pH 6.0 or pH 7.3, with 2 mM nitrate supplied. Cultures were grown under anaerobic conditions for 72 h and passaged 1:8 into fresh media for a total of four growth cycles. 16S amplicon sequencing was used to infer PD Nar^+^ relative abundance at the end of each cycle in pH 6.0 conditions, and at the end of four cycles in pH 7.3. (**E**) PD Nar^+^ relative abundance dynamics at pH 6.0 are shown. *f*_*Nar*+,0_ = 0.03, 0.5, and 0.97 are highlighted by darker lines. After a transient decrease in the *f_Nar_*+,_0_ *>* 0.1 conditions, co-cultures approach a value of *f_Nar_*+ ≈ 0.60–0.75. (**F**) PD Nar^+^ relative abundance dynamics at pH 7.3 are shown. The relative abundance of PD Nar^+^ approached zero after four cycles in all conditions, indicating that it was competitively excluded by RH Nap^+^ in this pH condition. All relative abundance values and uncertainties are calculated as described in Methods.

We conclude that the combination of PD Nar^+^ and RH Nap^+^ experienced substantially less toxicity due to nitrite than PD Nar^+^ by itself. We note that this community effect appears to derive from the apparent specializations of the two isolates on the resources nitrate and nitrite: PD Nar^+^ exhibited fast nitrate reduction and slow nitrite reduction, while RH Nap^+^ showed the opposite behavior (dark purple traces, Fig. 4A). While the nitrate specialization of PD Nar^+^ causes toxicity because it results in nitrite accumulation, the nitrite specialization of RH Nap^+^ allows it to remediate this toxicity.

We next turned to demonstrating that diminished nitrite toxicity in co-cultures of PD Nar^+^ and RH Nap^+^ explains the long-term dominance of PD Nar^+^ across multiple cycles of growth and dilution (Fig. 2B). To do this, we co-cultured these isolates over four cycles of serial dilution and measured relative abundances via 16S amplicon sequencing (Fig. 5D; Methods). In order to assess whether density-dependent effects determine the outcome of these experiments, we varied initial relative abundances of PD Nar^+^ and RH Nap^+^ over a wide range, holding total biomass constant.

Indeed, we found that after four cycles at pH 6.0, co-cultures across a range of initial relative abundances approached a stable configuration with PD Nar^+^ dominating (Fig. 5E). After the first cycle of growth, relative abundances of PD Nar^+^ decreased across most starting conditions, an effect perhaps attributable to physiological adaptation to anaerobic conditions (*48*). However, all conditions subsequently approached a state with PD Nar^+^ at 60–75% relative abundance by the fourth cycle. The amount of nitrite accumulation in these experiments was increased with the relative abundance of PD Nar^+^, with low abundances of PD Nar^+^ showing very little nitrite accumulation (orange/pink traces, Fig. S8G), and vice versa (blue/purple traces, Fig. S8G). This, along with the earlier results showing that RH Nap^+^ alleviates nitrite toxicity, supports the interpretation that PD Nar^+^ requires the presence of RH Nap^+^ at pH 6.0 to dominate.

Similar co-cultures performed at pH 7.3 showed RH Nap^+^ driving PD Nar^+^ to extinction after four cycles (Fig. 5F), corroborating our expectations from earlier experiments and analyses that RH Nap^+^ is simply a better competitor for resources at this pH condition (Fig. 3C, E, G; Fig. S7).

### Nar**^+^** and Nap**^+^** phenotypes are conserved across diverse taxa

Using isolates from our soil enrichment experiments (Fig. 2D), we demonstrated (Figs. 3-5) that the dominance of a Nar^+^ genotype (PD Nar^+^) under acidic conditions required the presence of a Nap^+^ genotype (RH Nap^+^). Underlying the interactions between these isolates was the fact that the former is an apparent nitrate specialist, capable of quickly consuming the primary limiting resource supplied to the community, while the later is a nitrite specialist, which mitigates the toxic effects of accumulated nitrite. Generalizing this observation, we next hypothesized that patterns of genotypes in the global topsoil microbiome (Fig. 1) arose due to similar interactions driven by the same patterns of specialization: namely, that Nar^+^ and Nap^+^ display nitrate and nitrite specialist phenotypes, respectively, meaning that the former genotypes tend to accumulate nitrite while the latter alleviate its toxicity.

First, we asked whether or not it is typical for denitrifiers to possess either Nar or Nap independently, rather than possessing both. An analysis of over 2000 denitrifier genomes in the KEGG reference genome database confirmed that this is indeed the case (Supplementary Information), showing a negative correlation between the presence/absence of these two genes (Fig. S9).

Second, we asked whether the traits and resource specializations of these genotypes are conserved across taxa. Recent work showed that the rates of nitrate and nitrite utilization by diverse denitrifying bacteria are strongly correlated with the presence and absence of denitrification genes they possess (*23*). As a result, we expect that Nar^+^ and Nap^+^ genotypes generically exhibit the phenotypes shown in Fig. 3.

To answer this question, we leveraged the library of phylogenetically diverse Nar^+^ and Nap^+^ strains isolated in our previous study (*23*). We identified 7 representative strains possessing one or the other nitrate reductase (3 Nar^+^, 4 Nap^+^) spanning *α*, *β* and *γ*-proteobacteria. To avoid ambiguity, we avoided strains with both Nar and Nap, although in such strains Nap is likely used for redox balancing (*49*). For each strain, we measured nitrate and nitrite utilization phenotypes at pH 6.0 supplied with 1.75 mM nitrate, using the area under the curve (AUC) of nitrate and nitrite dynamics to summarize traits (Fig. 6A). Values of nitrate AUC are inversely proportional to the rate of nitrate consumption (top panel, Fig. 6A), while values of nitrite AUC are directly proportional to nitrite accumulation (bottom panel, Fig. 6A).

**Figure 6:**
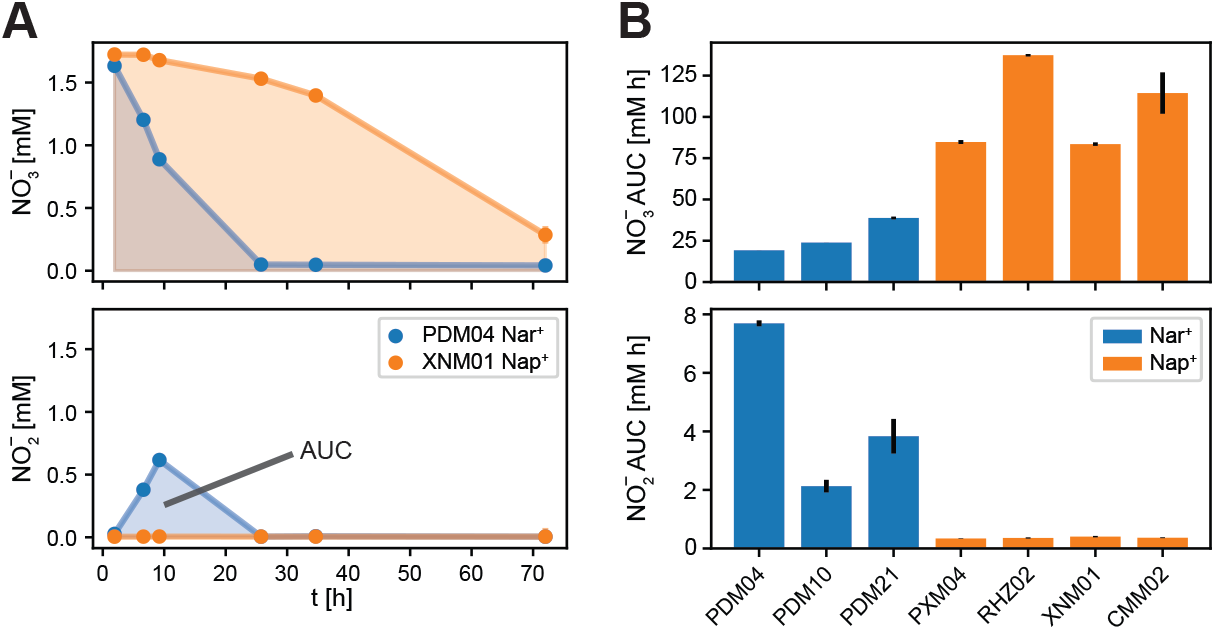
Nar^+^ and Nap^+^ phenotypes are conserved across diverse taxa. Additional Nar^+^ and Nap^+^ soil isolates were grown in anaerobic conditions at pH 6.0, and their denitrification phenotypes were measured. (**A**) Denitrification phenotypes were summarized by area under the nitrate (top) and nitrite (bottom) curves (AUC). Metabolite dynamics for PDM04 Nar^+^ (blue) and XNM01 Nap^+^ (orange) monocultures are shown, with area under the curve (AUC) for each metabolite illustrated by the colored shaded regions. Smaller AUC indicates faster metabolite reduction. (**B**) AUC for nitrate (top) and nitrite (bottom) for three Nar^+^ and four Nap^+^ strains isolated in Ref. 23. Nar^+^ strains exhibit faster nitrate reduction (blue bars, top panel) and slower nitrite reduction (blue bars, bottom panel), and Nap^+^ strains are the opposite. The mean and standard deviation of the AUC across technical replicates (*n* = 2) are shown.

As with PD Nar^+^ and RH Nap^+^, we found that Nar^+^ genotypes all exhibited fast nitrate reduction and the transient accumulation of nitrite (blue bars, Fig. 6B). In contrast, Nap^+^ strains all exhibited slow nitrate reduction and no nitrite accumulation (orange bars, Fig. 6B). We conclude that Nar^+^ genotypes generically accumulate nitrite via fast nitrate reduction, and that Nap^+^ strains generically avoid this accumulation. This observation lends support to the idea that global patterns in gene content emerge from regularity in the interactions between genotypes due to conserved genotype-phenotype relationships.

### Proteobacteria dominate denitrifying populations in topsoils

The taxa characterized in Fig. 6 are restricted to *α*, *β* and *γ-*proteobacteria, suggesting that the genotype-resource specialization traits we observed in PD Nar^+^ and RH Nap^+^ are likely conserved within the phylum Proteobacteria. However, soil microbiomes harbor substantial taxonomic diversity, raising the question of whether or not the trait conservation we observe in Proteobacteria is relevant to the high diversity communities present in the wild.

Therefore, we asked whether or not Proteobacteria were representative of the dominant denitrifying populations in soils. First, we performed a taxonomic analysis of the metagenomic reads annotated as denitrification reductases in the global topsoil microbiome (Fig. 1). We found that Proteobacteria indeed dominated these reads (Fig. S10), suggesting that a substantial portion of taxa possessing denitrification genes in topsoils are Proteobacteria. Second, we performed a detailed meta-analysis of existing literature examining the taxonomy of denitrifying populations in soils. We examined four studies based on genomic or taxonomic analyses of isolates from topsoils across diverse soil-types, pH, and geographic locations (Table S3). Ubiquitiously, we found that Proteobacteria dominated culturable denitrifying populations in soils, in many cases comprising *>*90% of the culturable population, again suggesting the dominance of Proteobacteria. However, since these studies were based on culturing isolates the dominance of Proteobacteria may reflect culture bias and not the true taxonomic diversity of denitrifiers in soils.

In order to overcome difficulties associated with culture bias, we performed an experiment to assess the taxonomy of denitrifying communities in natural soils using culture-free methods. We first collected 10 top soils across a range of pH values (5.0–7.1, Table S4). We brought these soils into the lab and following established methods created two types of microcosms, with (nitrate^+^) and without (control) nitrate amendments. Soils were incubated in anaerobic conditions for four days, nitrate utilization was measured and beginning and endpoint populations were sequenced to assess taxonomic structure (16S amplicon, Methods). By comparing ASV abundances after incubation in nitrate^+^ and control conditions, we quantitatively determined the taxa that respond to the presence of nitrate, while controlling for other metabolic processes such as fermentation. ASVs identified in this way are shown in Fig. 7, and again we find that Proteobacteria dominate nitrate-utilizing populations in top soils (Fig. 7B).

**Figure 7:**
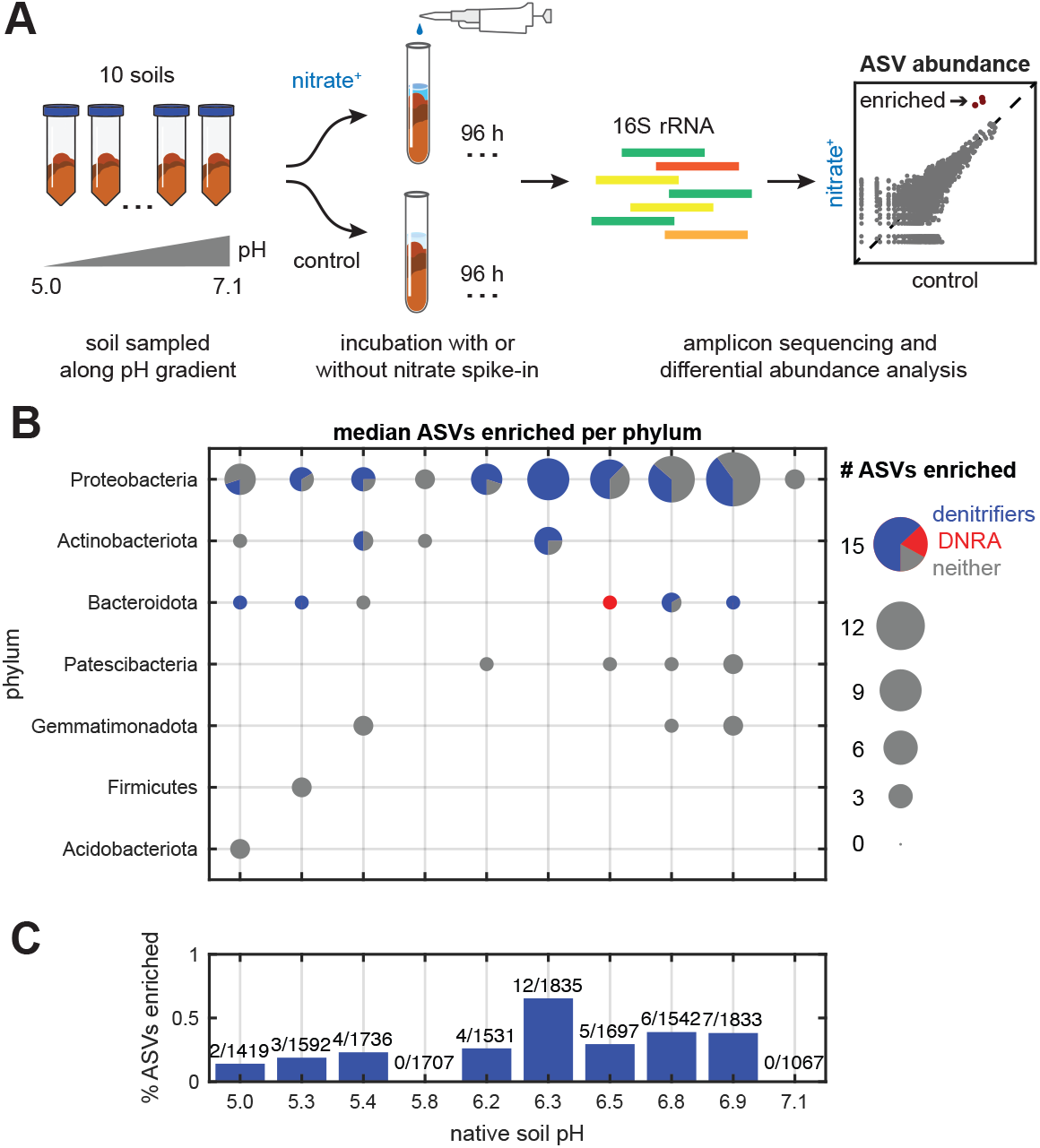
Proteobacteria dominate low-richness denitrifying populations in topsoils. Soil incubation experiments were performed to assess the response of endogenous communities to nitrate spike-ins. **(A)** Ten soil samples spanning a range of pH values (5.0–7.1) were collected from sites within the Cook Agronomy Farm (Pullman, WA, USA). Samples were processed to obtain soil slurries, which were then incubated in triplicate under anaerobic conditions for 96 h with spike-ins of nitrate which yielded 2 mM final concentrations (nitrate^+^) and without (control). DNA was extracted from frozen endpoint samples for 16S V3-V4 amplicon sequencing, and differential abundance analysis of amplicon sequence variants (ASVs) was performed to statistically determine which taxa were significantly enriched under addition of nitrate (red points, scatter plot). Details describing these procedures are given in Methods. **(B)** The number of ASVs (grouped by phylum) significantly enriched by nitrate spike-ins are shown for each soil (ordered by increasing pH). Pie charts show the fraction of ASVs classified by PICRUSt2 (*50*) as possessing the enzymes to perform denitrification (blue), DNRA (red), or neither (grey). Median ASV counts and genotype fractions measured across technical replicate incubations (*N* = 3) are shown. Across the range of pH values, Proteobacterial ASVs are most frequently enriched, with a substantial fraction of these ASVs likely possessing the genes to perform denitrification. **(C)** Bars show the percentage of enriched denitrifying ASVs out of total detected ASVs across the range of soils, and fractions are shown above the bars. Median values across technical replicate incubations (*N* = 3) are given. The small fraction of significantly enriched denitrifying ASVs suggests that denitrifying communities in topsoils are low-richness.

### Denitrifying communities in topsoils are low-richness

Our analysis of the interaction between PD Nar^+^ and RH Nap^+^ focuses on just two taxa. However, soils support thousands of taxa, raising the question of how an interaction between two taxa can be relevant in the complex ecological context of soils. However, our soil microcosm experiment provides strong evidence that the complexity of the nitrate-utilizing community in soils is low (Fig. 7). Specifically, a small fraction of all taxa in the soil community respond to the addition of nitrate (Fig. 7C). The result suggests that despite the ecological complexity of soils, just a handful of taxa participate significantly in this metabolic process *in situ*. Thus, interactions between a small number of taxa are potentially relevant for this metabolic process in soils, further supporting the idea that the interaction between Nar^+^ and Nap^+^ genotypes is relevant to patterns across the global topsoil microbiome.

## Discussion

Surveys of microbiomes in contexts from hosts to oceans have elucidated environmentally-mediated patterns in the composition of these complex communities. Understanding these patterns, and ultimately their implications for the function and persistence of microbial consortia is a long-standing problem in ecology. We have presented a data-driven and empirically validated approach to un-derstanding how changing environmental conditions drive changing community composition at the level of gene content. Our results reveal that environmentally mediated patterns on a global scale can emerge from interactions between members of a community. We arrived at this conclusion by first dissecting which environmental factors correlate with variation in gene abundances in the global topsoil microbiome, finding pH to be strongly correlated with the composition of the denitrification pathway. Our statistical approach motivated an experiment varying pH and assaying the structure of resulting denitrifying communities in the laboratory. Our enrichments recapitulated the observed pattern, allowing us to dissect interactions quantitatively. We showed that the dominance of the Nar^+^ genotype in acidic conditions relies on the presence of a Nap+ genotype to reduce pH-dependent nitrite toxicity.

The central finding of our study is that patterns in the relative abundances of genes on a global scale can emerge from ecological interactions between members of the community. The finding points to the key idea that patterns can emerge from interactions in communities as a whole, and not necessarily from specific traits conferring an advantage to individual taxa in certain environmental conditions (*10*). While filtering clearly plays a role in defining these patterns, for example at neutral pH in our study, interactions are likely essential to understanding the existence of patterns on a global scale.

### Limitations and extensions

Our analysis of gene content in the global topsoil microbiome revealed a pattern of gene abundances across a gradient of pH from very acidic soils (pH ∼4) to alkaline (pH ∼8). Over this range, we observed continuous variation in the abundances of the narG and napA genes. However, our monoculture and coculture experiments were restricted to pH 6.0 and 7.3. As a result, it remains unclear whether or not other interactions, or even filtering, might play a role in defining the full extent of the global pattern we observe. For example, it is common to find obligate nitrate reducers possessing *narG* that do not reduce nitrite (*23*). Additionally, we note that in enrichments at pH *<* 5.5, we do not find taxa of the Nap genotype, only Nar (Supplemental Information). Therefore, it may be that additional ecological and physiological processes are important in more acidic conditions. Interactions involving such genotypes and environments could be essential for fully understanding the global pattern we observed. Interrogating interactions in more acidic conditions could be useful in uncovering additional processes.

Next, while we provide strong evidence of the generality of the observed interaction, we do not show direct evidence of the toxicity-alleviation interaction between PD Nar^+^ and RH Nap^+^ in other strains. Thus, this interaction may not be present between all pairs of Nar^+^ and Nap^+^ genotypes at every pH. We show via simulation, however, that such interactions are likely to arise from ecological interactions that give rise to complementary metabolic phenotypes (Figs. 5 S13).

Finally, our statistical analysis of variation in pathway magnitude and composition focused on variations in the composition of the denitrification pathway across the globe. As a result, our study shows that interactions are important for determining which genotypes participate in the process of denitrification. However, the magnitude of the pathway, i.e., approximately the fraction of reads in the metagenome mapping to denitrification enzymes, decreased with increasing C/N ratio in soils (*ρ<* 0, Fig. S1). Denitrification is known to compete with dissimilatory nitrate reduction to ammonia (DNRA) in anaerobic environments (*31*), with low C/N favoring denitrification over DNRA for reasons of redox balance (*51*). Our findings are consistent since rising C/N inhibits denitrification globally. In this sense, it is likely that an environmental filtering framework does indeed explain which process, denitrification versus DNRA, dominates in a given environmental context. It will be an important avenue for future work to use the methods developed here to generate concrete hypotheses for the role of filtering and interactions in pattern formation. Testing these will be important for understanding the fate of nutrients in the environment, as DNRA results in the production of bioavailable ammonia, and denitrification drives nitrogen loss to N_2_.

### Trait conservation and trade-offs

A key finding of our study is the idea that consistent patterns in gene content on a global scale can reflect conserved genotype-phenotype relationships across diverse bacteria. This result is in agreement with our previous work showing that denitrification genotypes confer similar phenotypes across different taxa (*23*). The present study suggests that this conservation of traits can lead to the conservation of interactions between genotypes and thus patterns observable via metagenomic sequencing over many environments.

In our system, the interaction between Nar^+^ and Nap^+^ genotypes arises due to specialization by these two genotypes into the first and second steps of the denitrification cascade. This specialization suggests the possibility that there is a physiological trade-off between nitrate and nitrite reduction rates. Previous studies have implicated such a trade-off and suggested that it might emerge from competition for intracellular resources (*33, 52*). We investigated this further using simulations that account for the trade-off between nitrate and nitrite reduction rates and the toxicity of nitrite, which preferentially inhibits nitrate specialists (Supplemental Information). These simulations identified two phases of community assembly: (I) with high toxicity (low pH) and concave trade-off, we observed a regime where two generalist strains that perform both nitrate and nitrite reduction coexist (Fig. S12B), and (II) when toxicity is low (neutral pH) and the front is concave, a single generalist (reduces both nitrate and nitrite) dominates (Fig. S12B). Thus, we found that the observed experimental phenomena at both pH 6.0 (regime I) and pH 7.3 (regime II) emerge generically from the imposition of empirically observed physiological constraints. Together, these findings suggest the idea that learning conserved physiological constraints on traits will be an important step in understanding ecological processes and patterns on a global scale.

### The role of enzyme properties in denitrification traits

In this work we have used the genes *narG* and *napA* as genetic markers of conserved, interacting, phenotypes where strains with *narG* are nitrate specialists and strains with *napA* are nitrite specialists. Here we discuss how the enzymes that make up the pathways might mechanistically confer the traits we observe. Consistent with our results here we previously showed that nitrate reducers possessing *narG* instead of *napA* exhibit faster nitrate reduction (*44*), a finding supported by *in vitro* studies of these enzymes (*53–57*). In addition, the Nar nitrate reductase is protected from extracellular pH in the cytoplasm, while the periplasmic Nap reductases are not (*24*), possibly explaining faster nitrate reduction at low pH for Nar^+^ strains.

A key finding of our study is the alleviation of nitrite toxicity, and thus the nitrite reductases encoded by the genes *nirS* and *nirK* are also likely important. Statistically, we observe *nirS* and *narG* have the same relationship with pH as do *napA* and *nirK*. This observation aligns with the fact that nitrite toxicity alleviation is accomplished by a denitrifier possessing *nirK* (Fig. 2), suggesting that nitrite specialists possess the NirK nitrite reductase and not the NirS enzyme (Figs. 1H, 2D; additionally, every Nar^+^ strain in Fig. 6B also contains *NirS* except PDM21, and every Nap^+^ strain also contains *NirK*). This is also consistent with previous work, which shows NirK enzymes have higher activity than NirS (*24, 58–71*), and the presence of *nirK* rather than *nirS* predicts a faster nitrite reduction traits (*44*). Thus, qualitatively the known properties of enzymes in the pathway accord with the nitrate and nitrite specialization we observe in our isolates. In addition, studies examining the transcriptional and translational dynamics of reductases of denitrifiers to changing pH show that *nirS* possessing strains have decreased enzyme levels at low pH (*72, 73*) while NirK possessing strains do not (*74–76*). A search of the BRENDA (*77*) database for experiments on purified NirS and NirK enzymes did not show significant differences in pH optima (*78–84*). Thus, the nitrite reductases likely have differential basal activity and pH alters their transcriptional or translational dynamics. Together, the properties of the denitrification pathway qualitatively agree with the traits we observe for these genotypes.

### Learning from environmental variation in the wild

Our results demonstrate the power of statistical analysis to motivate *in vitro* experiments that connect strain-level phenomena to macroecological patterns. Our approach used a statistical decomposition of variation in natural communities to reveal patterns in gene abundances. This approach is in contrast to a “mechanism-first” approach recently undertaken by Abreu *et al.* (*85*) that identified the phenomenon of slow growth being favored at warmer temperatures in a simplified laboratory community, before searching for this pattern in the wild. While both approaches are powerful for understanding patterns in nature, a statistical approach allows us to discover patterns and processes without first postulating the environmental variables responsible or the details of the interaction in the laboratory. In this way, our approach allows us to uncover the key environmental drivers of variation in the wild before dissecting the role of interactions in the community.

### The stress gradient hypothesis

The stress gradient hypothesis (SGH) provides a framework for understanding how the sign of ecological interactions depends on environmental conditions (*86*). The SGH suggests that interactions should skew positive (facilitation) in the presence of an environmental stressor, and negative (competitive) in the absence of stress. Previous attempts to demonstrate the SGH in the plant and microbial world highlight challenges associated with the ambiguity of the hypothesis (*87*). Namely, not all stressors appear to select for facilitative interactions (*88*), and in situations where the SGH has been observed (*89*), it is unclear whether any conserved underlying mechanisms are responsible. These challenges mean that we do not yet know whether a systematic description of the impact of stress on community interactions is plausible.

Our results represent a clear demonstration of the SGH, with nitrite toxicity at low pH serving as the stressor and the alleviation of toxicity by Nap^+^ genotypes serving as the facilitative interaction. More importantly, our approach demonstrates a systematic methodology for elucidating the SGH in communities. Specifically, we have shown that conserved genotype-phenotype relationships can drive patterns of facilitative interactions, leaving a signature of the SGH in covariation between environmental variables and metagenomic data. Further analyses of these datasets could yield insights into the importance of the SGH for other microbial processes. Such analyses could enable a more systematic interrogation of the relevance of the SGH in structuring natural communities.

### The environmental filtering hypothesis

The concept of environmental filtering has a long history, perhaps beginning with the foundational work of von Humbolt and Bonpland (*90*), and the belief that traits alone can determine community composition has remained persistent when interpreting data in the modern era. Surveys in marine microbiomes show that functional traits correlate with environmental variables (*91*). Similarly, studies of the gut microbiome have interpreted reproducible changes in the abundances of specific taxa in response to diet in terms of environmental filtering (*92*). In these very complex ecological contexts, it is challenging to definitively determine when interactions matter and when they do not. Experimental and analytical methods for dissecting interactions in these contexts will be important going forward.

More broadly, we hope this work helps establish a link between two conceptual frameworks in community ecology. First, decades of work have focused on the structure of interactions between taxa and the resulting abundance dynamics (*93*). In this picture, traits are often assigned randomly, and community function or ecosystem services play a secondary role. Second, quantitative physiology and an appreciation of the importance of metabolic traits in community assembly (*94*) have revealed that traits are not random, but rather that strong conservation yields regular structure in the physiology of microbes, yielding robust quantitative principles (e.g., Ref. 95). Our study proposes a path forward that bridges these two conceptual frameworks, revealing how conserved traits can give rise to recurrent patterns of interactions in wild communities with implications for ecosystem functions. We hope that understanding the origins of these patterns will drive deeper insights into the assembly and function of complex natural microbiomes.

## Acknowledgments

We acknowledge Luis Miguel de Jesús Astacio for collecting soils, Rita Almeida Oliveira for help with colony counting technique, Mohammad Bahram for previously published global topsoil microbiome data, and Kabir Husain for useful discussions. Soil sampling done in the Cook Agronomy Farm was, in part, a contribution from the Long-Term Agroecosystem Research (LTAR) network. LTAR is supported by the United States Department of Agriculture.

## Author contributions

K.C., K.G., M.M. and S.K. conceptualized the research. K.C., K.G., and S.K. designed the experiments. K.C., K.G., K.L., and M.C-W. performed the experiments. K.G. performed statistical analysis of the topsoil microbiome data and performed and analyzed soil enrichment experiments. M.C-W. performed mortality rate experiments, advised by K.C. and S.K.. Z.L. performed co-evolution analysis, advised by K.G. and S.K.. K.C. performed all other experiments and analyses, advised by K.G. and S.K.. K.C. performed CRM fits, predictions, and simulations, adapting code written by K.G.. K.L. sampled topsoils and performed the soil microcosm experiments. M.T. performed the analysis of sequencing data for the soil microcosm experiment in Fig. 7. The topsoil microcosm experiment was, in part, a contribution from the Long-Term Agroecosystem Research (LTAR) network. LTAR is supported by the United States Department of Agriculture. K.C., K.G., Z.L., K.L., M.T., and S.K. wrote the manuscript with input from all authors.

## Data and materials availability

Code and data associated with this manuscript are publicly available at https://github.com/kyle-crocker/interaction-patterns and by request. Raw sequence reads for enrichment experiments are deposited under NCBI BioProject ID PRJNA976277. Bacterial isolates are available by request.

