## Supplementary material for "Global patterns in gene content of soil microbiomes emerge from microbial interactions": Methods and Supplementary Information

#### Analysis of the global topsoil microbiome

##### Data pre-processing

Gene abundance tables and environmental variables for the global topsoil microbiome dataset were provided by Bahram *et al.* (*3*), who sampled, sequenced, and analyzed 189 topsoil sites representing the world's terrestrial biomes. Samples were processed according to standardized protocols, including shotgun metagenomic sequencing of soil DNA extracts, and chemical analysis of soil pH, P, K, Ca, Mg, $^{12}C$, $^{13}C$, $^{14}N$, and $^{15}N$. In addition, site data including mean annual temperature (MAT), mean annual precipitation (MAP), potential evapotranspiration (PET), net primary productivity (NPP), and moisture were obtained from public databases. Paired metagenome reads were quality filtered and annotated to yield KEGG ortholog (KO) abundance tables. The details of all protocols and analyses to obtain gene abundance tables and site environmental variables are given in Ref. 3.

In the subsequent analyses, the abundance of KOs corresponding to the six reductases in the denitrification pathway (*narG*/K00370, *napA*/K02567, *nirS*/K15864, *nirK*/K00368, *norB*/K04561, *nosZ*/K00376) were considered. The relative abundances of these KOs were computed by dividing the number of reads mapped to each KO by the total number of reads in each sample. In addition, 17 site environmental variables from the measurements and analyses described above were considered: pH, P, K, Ca, Mg, C, N, $\delta^{13}C$, $\delta^{15}N$, C/N, latitude, longitude, MAT, MAP, PET, NPP, and moisture.

##### Unit-invariant singular value decomposition

Unit-invariant singular value decomposition (uiSVD; Ref. 30) was applied to the denitrification reductase relative abundance matrix ($X$) to separate variation in pathway magnitude (i.e., variations in total reductase content) from variation in pathway composition (i.e., variations in relative fractions of reductase content). This approach was motivated by modeling the data as

$$\vec{x}_i = d_i \vec{c}_i, \tag{1}$$

where $\vec{x}_i$ is the $i$th row of $X$ (with entries being relative abundances of each KO), $d_i$ is a scalar representing pathway magnitude and $\vec{c}_i$ is a vector representing pathway composition. In matrix form, this can be written as

$$X = DC, \tag{2}$$

where $D$ is a diagonal matrix. Expressing $X$ in this form can be achieved by applying uiSVD, which decomposes $X$ as

$$X = D\tilde{U}\tilde{S}\tilde{V}^T E, \tag{3}$$

where $D$, $\tilde{S}$, and $E$ are diagonal matrices and $\tilde{U}$ and $\tilde{V}$ are unitary matrices. Hence

$$C = \tilde{U}\tilde{S}\tilde{V}^T E. \tag{4}$$

The results of this decomposition are shown in Fig. 1D. Note that, in this representation, $D$ encodes the scale of the rows of $X$ (i.e., pathway magnitude), while $E$ encodes the relative scales of the columns (i.e., typical relative scales of each gene). Code used to perform uiSVD was adapted for Python from the MATLAB code given in (*30*).

Having decomposed the relative abundance data $X$ into elements of pathway magnitude ($D$) and composition ($C$), modes of gene covariation within the composition matrix were identified via principal components analysis (PCA). Because $E$ encodes scaling information about the genes in $X$, $\tilde{C} \equiv CE^{-1}$ can be interpreted as a normalized gene composition matrix, for which all genes have been set to the same approximate scale. Therefore PCA was applied to $\tilde{C}$ by mean-centering the columns, and applying singular value decomposition the resulting matrix, yielding a decomposition $USV^T$. The columns of $V$ represent the orthonormal principal components (PCs) of $\tilde{C}$ (shown in Fig. S2), and

$$T = US \tag{5}$$

represents the "scores" matrix, i.e., the projections of $\tilde{C}$ onto the PCs in $V$.

#### On the compositional nature of shotgun metagenomic data

While relative abundance data are, strictly-speaking, subject to the effects of compositionality (*96*), the conventional log-ratio transforms attributable to Aitchison (*97*) *were not* applied to the data prior to decomposition via uiSVD. This choice was motivated by the substantial challenges associated with the treatment of zeros in the data prior to transformation, which cause the logarithm to diverge to $-\infty$. While strategies have been devised for imputing zeros in some circumstances (e.g., Refs. 98, 99), robust imputation necessarily relies on estimating underlying distributions for sequencing read counts, using for instance data from multiple samples from an experimental group or condition. Arguably such strategies cannot be sensibly applied in the present study, since doing so would require *a priori* assuming underlying similarities between the samples which indeed we are attempting to infer through our analyses.

The choice to forgo log-ratio transformation of the denitrification reductase relative abundances from the global topsoil microbiome dataset is justifiable in the following way. In short, it is possible to show that the effects of compositionality (in particular the "negative bias" associated with correlations, Ref. 96) are negligible for these data. At an intuitive level, this arises because the denitrification reductases make up a small fraction (typically no more than one part in 10,000) of the gene sequences in the dataset. To see this first consider the simplest case of a relative abundance dataset with only two variables, i.e., two genes or taxa. The spurious effects of compositionality on correlations in this case are maximal; an increase in one variable *must* correspond to a decrease in the other, even in the scenario where the absolute abundances of both variables simultaneously increase, with one simply increasing more than the other. In the opposite limit of many variables, with no one in particular making up a substantial fraction of the composition, an increase in the relative abundance of one variable need not result in a substantial change in any other variable, because changes in this variable are small relative to what is described in the first scenario, and there are many other variables to "absorb" the small change. This phenomenon has been noted before, for instance in Ref. 100, wherein Friedman and Alm examined the effect of sample diversity on the strength of compositional effects, as measured by the fidelity of correlations inferred on relative abundance data. The authors found that dependencies between taxa in synthetic datasets were accurately inferred using naive Pearson correlations in the case of high diversity datasets. The diversity in these datasets corresponded to a Shannon number of effective OTUs (exponential Shannon entropy) $n_{eff} = 30$. For comparison, the corresponding median value across all 189 samples and 7700 KOs for the full global topsoil microbiome dataset is $n_{eff} = 1837$, which clearly represents an even greater level of diversity. Similar results relating sample diversity to the fidelity of recovered correlations have been found in a more systematic comparison of approaches (*101*).

This intuition can be made more formal by showing that the constituent operations of Aitchison's transforms, a logarithm and a ratio, are often not necessary to eliminate negative bias in high-diversity datasets. Negatively-biased correlations in compositional data are a consequence of the unit-sum constraint, i.e., that the values of relative abundances in a sample must sum to one. Aitchison's transforms are conventionally used to eliminate this constraint through two operations: by first taking a ratio, and then taking the logarithm. In the case of the additive log-ratio, an $N$-dimensional compositional vector $\vec{x} = (x_1, x_2, ..., x_N)$ is transformed to an $N-1$-dimensional vector by choosing a variable for taking the ratio, e.g., $\text{alr}(\vec{x}) = (\log(x_1/x_N), \log(x_2/x_N), ..., \log(x_{N-1}/x_N))$, where $x_N$ here has been chosen for the ratio. However, Rayens and Srinivasan (*102*) point out that it is the ratio and not the logarithm in Aitchison's transforms that eliminate the unit-sum constraint. That is, it can be proven that negative bias in correlations is eliminated after the ratio operation, and that taking a logarithm is superfluous to this goal[1]. Going further, even the ratio is often not strictly necessary. It is frequently the case in high-dimensional and high-diversity datasets that at least one variable can be identified for taking the ratio which, by chance, is approximately constant across samples (*103*). Since dividing variables by an approximately constant value has little influence on the values of correlations between those variables, taking a ratio in this scenario is superfluous to eliminating negative bias as well. In the case of the denitrification reductase relative abundances in

[1]Rayens and Srinivasan also show that the logarithm serves primarily to transform the ratioed data to normality, and can be replaced with other Box-Cox transforms in the pursuit of this end.

the global topsoil microbiome, the sum total of all other gene variables can be taken as a convenient choice for demonstrating that taking a ratio is unnecessary. By defining a 7-variable compositional vector, where the first six variables are the denitrification gene relative abundances and the seventh is the total relative abundance of all other genes, a ratio can be defined between the denitrification genes and the total of the other genes. Since the total relative abundance of other genes ranges between 0.9998 and 1.0, the original relative abundances themselves are a very good approximation for the ratio.

It is worth noting that the arguments above are generally not specific to the global topsoil gene dataset, and would likely apply to most other gene relative abundance datasets obtained via shotgun metagenomic sequencing, owing to the likelihood of high effective diversity in such datasets.

#### Analyzing gene-environment covariation

In order to relate gene covariation to variation in site environmental variables, Pearson correlations ($\rho^2$) were computed between the scores of each principal component (Eq. 5) and each environmental variable (squared Pearson correlations shown in Fig. 1F). Additionally, the same analysis was performed between the diagonal elements of $D$ and each environmental parameter (Fig. 1E). Each correlation was computed using the $n = 189$ observations from the global topsoil microbiome dataset, with environmental variables containing NaN values removed when computing each correlation. The significance of associations between C/N ratio and $D$, as well as between pH and PC2 scores was determined via a one-tailed randomization test. Explicitly, a distribution for the null hypothesis of zero correlation was empirically constructed by repeatedly ($10^6$ times) computing correlations between shuffled versions of the variables.

#### Phylogenetic classification of denitrification reductases

Raw reads from the global topsoil microbiome dataset (PRJEB18701) were trimmed of adapters and low quality sequences using Trimmomatic ver. 0.39 with default settings. Trimmed reads were assembled using SPAades ver. 3.15.0. Predicted ORFs on assembled contigs were then annotated as KEGG ortholog groups (KOs) of the denitrification reductases (*narG*/K00370, *napA*/K02567, *nirS*/K15864, *nirK*/K00368, *norB*/K04561, *nosZ*/K00376) using eggNOG-mapper ver. 2.0.8, and reads were mapped to these annotated ORFs using minimap2. Finally, reads mapping to denitrification KOs were phylogenetically classified using the Kaiju web server (`https://kaiju.binf.ku.dk/server`; Ref. 104). The results of these classifications, at the level of taxonomic phylum and grouping all samples together, are shown in Fig. S10.

#### Enrichment and isolation of denitrifying strains along a pH gradient

##### Processing of soils for primary enrichment experiments

Six forest and prairie soil samples were collected from Meadowbrook Park, Urbana, IL. Details regarding these soil samples are given in Table S1. Soils were sampled from a depth of 1–5 cm using autoclaved steel laboratory spatulas, and then stored in sealed plastic bags at 4 °C for approximately two months prior to the start of the experiment. No sample compositing was performed. To mechanically homogenize soils prior to enrichment, 5 g samples of each soil were added to sterile 50 mL centrifuge tubes, along with 25 mL PBS (pH 7.4) and 5–10 g sterile 4 mm glass beads. Tubes were then vortexed (Vortex-Genie 2) at high speed for 1 min to homogenize the samples. After vortexing, large particles in homogenized soil samples were allowed to settle for 20 min before transferring 1 mL of supernatant to sterile microcentrifuge tubes.

##### Primary enrichment experiments

Wells of a sterile 96-deepwell plate (Axygen PDW20C) were loaded with 1.2 mL of defined media. The media contained succinate (25 mM) as the carbon source and 2.0 mM sodium nitrate was initially supplied. Medium pH was buffered by phosphate (40 mM) at two conditions, pH 6.0 and 7.3. This medium will hereafter be referred to as succinate defined medium (SDM); its precise composition is described in Ref. 23, and it was developed to capture a diverse range of denitrifiers (*26*). Six wells of each pH condition were then inoculated with 10 µL of each soil supernatant, with two wells of each pH condition left as no-growth controls. The plate was then sealed with a gas-permeable sterile membrane (Diversified Biotech BERM-2000). After sealing, the plate was immediately transferred to an anaerobic glove box (Coy Laboratory Products 7601-110/220), which was continuously purged by a 99%/1% $N_2$/$CO_2$ mixture. The plate was incubated at 30 °C and shaken at 950 RPM (Talboys Professional 1000MP, 3 mm orbital radius).

Cultures were grown under these conditions for 72 h. At the end of this time, 150 µL of the cultures were passaged under anaerobic conditions into a freshly prepared plate containing 1050 µL of fresh medium (1/8 dilution factor). These passaged cultures were sealed and returned to incubation and shaking for another 72 h growth cycle. At the end of each cycle, unused cultures were assayed for endpoint nitrate and nitrite concentrations via Griess assay and vanadium (III) chloride reduction method via the protocol described in Ref. 23 (Fig. S3). In addition, optical densities at 600 nm were also recorded (BMG CLARIOstar) using 300 µL of endpoint cultures in 96 well optical plates (Fig. S3).

Cultures were repeatedly passaged and assayed in this manner for 12 cycles. At the endpoints of cycles 4, 8, and 12, 100 µL samples of endpoint cultures were cryopreserved by adding 100 µL of 50 % glycerol and storing at −80 °C. At the end of cycle 12, cultures remaining after cryopreservation and nitrate/nitrite assay were frozen at −20 °C for subsequent DNA extraction.

**Sequencing and analysis of primary enrichment experiments**

Frozen enrichment endpoint (cycle 12) cultures stored at −20 °C were thawed and DNA extracted using the DNeasy UltraClean Microbial Kit (Qiagen). DNA concentrations were quantified using the Qubit dsDNA BR Assay Kit (Invitrogen). Library preparation for sequencing of DNA extracts was performed using the Nextera DNA Prep Kit and the Nextera DNA CD Indexes (Illumina). Pooled libraries were then sequenced using a NextSeq 500/550 Mid Output Kit v2.5 (Illumina, 2 × 150 bp paired-end), with a 1.5 pM library loading concentration and a 1% spike-in of PhiX Control v3 (Illumina). Sequencing was performed on a locally maintained and operated Illumina NextSeq 550 system.

To characterize the taxonomic composition of enrichments, 16S rRNA fragments (miTAGs) were extracted from the sequencing data using the miTAGs extraction script ver. 1 (*39*), after trimming and merging of overlapping paired-end reads via Trim-Galore ver. 0.6.7 (*105*) and BBMerge ver. 38.22 (*106*), respectively. The resulting miTAGs were then taxonomically classified using the RDP Classifier (*40*), using a confidence threshold of $80\%$ and the copy number adjustment option enabled.

The sequencing data were then binned into metagenome assembled genomes (MAGs) using the metaWRAP pipeline ver. 1.3 (*43*). In summary, the pipeline performed adapter and quality trimming of raw reads using Trim-Galore ver. 0.6.7 (*105*), assembly using metaSPAdes ver. 3.15.4 (*107*), binning using CONCOCT ver. 1.1.0 (*108*), MaxBin2 ver. 2.2.7 (*109*), and metaBAT ver. 2.15 (*110*), bin quality assessment using CheckM ver. 1.0.11 (*111*), and bin refinement by combining the results of the three binning algorithms. Bin completion and contamination thresholds of $95\%$ and $5\%$, respectively, were used to obtain high-quality MAGs in most samples (Table S2). High-quality MAGs were then annotated using the RAST server (*41*).

#### Isolation of strains from primary enrichment experiments

Strains representing the dominant taxa in the enrichment experiments were isolated from cryopreserved cycle $12$ samples from soil #1. Glycerol-cryopreserved cultures were streaked to purity on 1/10x TSB agar plates ($1.5\%$ agar w/v) in aerobic conditions. Overnight cultures of single-colony isolates were grown in 1/10x TSB ($30\,^{\circ}\mathrm{C}$, $400$ RPM) in aerobic conditions and cryopreserved ($50\%$ glycerol, $-80\,^{\circ}\mathrm{C}$). Sanger sequencing of the 16S rRNA gene using 27F and 806R universal primers was used to taxonomically classify isolates using the SILVA rRNA database (*112–114*). RH Nap$^{+}$ was determined to be of the family *Rhizobiaceae* and PD Nar$^{+}$ was determined to be of the genus *Pseudomonas*.

##### Additional enrichments across a broader pH range

Additional enrichments using a different set of soil samples were performed across a broader range of pH conditions, spanning pH 5.0 to 7.3. Seven soil samples (soils #7–13, Table S1) taken from prairies and forest preserves across the Midwestern United States (IL, IN, MI, WI) from a depth of 1–5 cm using autoclaved steel laboratory spatulas, and were homogenized and processed as described above. These soils were used to inoculate an enrichment experiment with four different pH-buffered SDM conditions: 5.0, 5.5, 6.0, 7.3. In pH 5.0 and 5.5 conditions, succinate/succinic acid serves as the buffering agent because the buffering capacity of phosphate is weak in this pH range. Details of incubation, passaging, sampling, cryopreservation, as well as of sequencing and data analysis of cycle 12 cultures, are as described above, except that defined media succinate concentration is reduced to 4 mM. Since nitrate is the limiting resource in these experiments, this change does not impact growth, but a lower succinate concentration reduces spontaneous protonation at low pH. The taxonomic composition and median MAG genotypes of endpoint enrichments are shown in Fig. S4.

#### Characterization of isolate phenotypes in denitrifying conditions

##### Culturing protocol

Strains were pre-cultured in two stages under aerobic conditions prior to transfer to denitrifying (anaerobic) conditions for phenotyping. First, wells of a sterile 24-well plate (Thermo Scientific Nunc Non-Treated Multidishes) were loaded with 1.7 mL of R2B medium. Wells were inoculated with isolates PD $Nar^+$ and RH $Nap^+$ from glycerol stocks stored at −80 °C. The plates were then sealed with a gas-permeable sterile membrane (Breathe-Easier, USA Scientific, 9126-2100). After sealing, the culture was incubated overnight at 0.5 rcf (400 RPM in Fisherbrand Incubating Microplate Shakers 02-217-759, 3 mm orbital radius or 219 RPM in Heidolph Unimax 1010, 10 mm orbital radius) and 30 °C in aerobic conditions. These cultures reached saturation during this time. Second, wells of a sterile 24-well plate were loaded with 1.7 mL of SDM at pH 7.3 with 25 mM succinate (and no sodium nitrate). Wells were then inoculated with $17\,\mu L$ of the saturated R2B PD Nar$^+$ and RH Nap$^+$. After sealing, the cultures were incubated at $0.5$ rcf and $30\,°C$ in aerobic conditions overnight. These cultures reached saturation during this time. Saturated SDM culture densities were measured and normalized to an optical density of $1.0$ (measured at $600\,nm$) via dilution into pH $7.4$ phosphate-buffered saline ($8\,g/L$ $H_2O$, $0.2\,g/L$ KCl, $2.68\,g/L$ $Na_2HPO_4{\cdot}7H_2O$, $0.24\,g/L$ $KH_2PO_4$).

Due to a lag time associated with growth in anaerobic conditions for facultative anaerobes (*115–120*), an additional period of pre-culture in anaerobic conditions was performed before phenotyping. Wells of a sterile $96$-deepwell plate (Axygen PDW20C) were loaded with $1.2\,mL$ SDM ($4\,mM$ succinate) supplemented with $1\,mM$ sodium nitrate, from stock that had been allowed to equilibrate in the anaerobic glovebox. SDM without nitrate was loaded into at least three wells of the plate as a control for growth on trace quantities of oxygen; since succinate is a non-fermentable carbon source, any growth in the absence of nitrate in this medium indicates aerobic growth. These wells were inoculated in the glovebox with $12\,\mu L$ of OD-normalized PD Nar$^+$ and RH Nap$^+$ aerobic pre-cultures, resulting in a starting OD of $0.01$. Additional wells were left blank as no-growth controls. Plates were sealed with a gas-permeable sterile membrane. Cultures were incubated at $30\,°C$ and shaken at $950\,RPM$ (Fisherbrand Incubating Microplate Shakers 02-217-759 or Talboys Professional 1000MP, 3 mm orbital radius) for $72\,h$.

Anaerobic pre-cultures were then used to initiate denitrification phenotyping experiments. Optical densities of endpoint anaerobic pre-cultures were measured using $300\,\mu L$ of cultures in $96$-well optical plates. Optical densities were then normalized to $0.04$ via dilution into SDM (no nitrate). $150\,\mu L$ of this normalized endpoint anaerobic culture was then passaged to $1050\,\mu L$ of SDM, with pH and nitrate/nitrite concentrations varying according to the experiment (e.g., Fig. 4). In the case of co-culture phenotyping experiments, this $150\,\mu L$ inoculum contained a 1:1 ratio of PD Nar$^+$ and RH Nap$^+$, as measured by optical density. Inoculated plates were then incubated at $30\,°C$ and shaken at $950\,rpm$ in anaerobic conditions. Nitrate and nitrite concentrations were assayed over time via manual sampling and subsequent Griess assay and vanadium (III) chloride reduction via the protocol described in Ref. 23. Endpoint biomass was measured via optical density at $600\,\mathrm{nm}$ using $300\,\mu\mathrm{L}$ samples.

Additional experiments were conducted where measurements were taken over multiple cycles of growth and dilution into fresh medium (e.g., Figs. 3 & 5). For these experiments, aerobic pre-cultures were used to directly inoculate the first cycle of anaerobic growth. Nitrate and nitrite measurements were taken beginning in the first cycle of growth, and normalization of optical density after the first cycle was omitted. For the first cycle, $1.2\,\mathrm{mL}$ of SDM ($4\,\mathrm{mM}$ succinate, $2\,\mathrm{mM}$ nitrate) was loaded into a 96-deepwell plate as described above. $12\,\mu\mathrm{L}$ of normalized culture from the endpoint of the aerobic pre-culture protocol was used to inoculate the wells of this plate. For the experiment shown in Fig. 5, the fractions of PD $\mathrm{Nar}^+$ and RH $\mathrm{Nap}^+$ were chosen to achieve specified relative abundances by optical density. Two no-growth controls that contain either no inoculum (to control for contamination) or no nitrate or nitrate (to control for aerobic growth) were included on each plate.

At the end of each growth experiment or cycle, cultures were removed from anaerobic conditions, sealed with Microseal 'B' Seals (BIO-RAD, MSB1001), and stored at $-80\,^{\circ}\mathrm{C}$ for subsequent DNA extraction.

**Quantification of relative abundance dynamics in co-culture**

Relative abundances were quantified in co-culture experiments using 16S amplicon sequencing. Frozen endpoint co-culture enrichments stored at $-80\,^{\circ}\mathrm{C}$ were thawed and DNA extracted using the DNeasy 96 Blood & Tissue Kit (Qiagen). DNA concentrations were quantified using the Qubit dsDNA BR Assay Kit (Invitrogen). Next, the 16S v4 region of the rRNA gene was amplified with 515F and 806R primers using the Illumina 16S sequencing protocol (*121*). Briefly, a fragment of the 16S rRNA gene was amplified using the 515F (GTGYCAGCMGCCGCGGTAA) and 806R (GGACTACNVGGGTWTCTAAT) universal primers. The following reagents were used for each reaction: $14.5\,\mu L$ nuclease-free $H_2O$, $1\,\mu L$ 515F primer ($5\,\mu M$), $1\,\mu L$ 806R primer ($5\,\mu M$), $10.5\,\mu L$ DNA extract, $12.5\,\mu L$ Platinum Hot Start PCR Master Mix (Invitrogen). The following thermocycler settings were used: initial denaturation, $3\,\mathrm{min}$ at $95\,°C$; amplification (25 cycles), $30\,\mathrm{sec}$ at $95\,°C$, $30\,\mathrm{sec}$ at $55\,°C$, $30\,\mathrm{sec}$ at $72\,°C$; final extension, $5\,\mathrm{min}$ at $72\,°C$. PCR products were cleaned using the PCRCLEAN DX Kit (Aline). 16S amplicons were barcoded for multiplexed Illumina sequencing using the Nextera XT Index kit (Illumina) following the Illumina protocol (*121*). Pooled amplicon libraries were then submitted to the University of Chicago Genomics Facility for sequencing using a 15-20% spike-in of PhiX Control.

Resulting paired-end reads were merged with Pear (*122*) and then quality filtered with DADA2 plugin in the QIIME2 pipeline with default parameters (*123*). Taxonomy of ASVs were assigned by q2-feature-classifier prefitted to the SILVA database using the naive Bayes algorithm for the V4 region of 16S rRNA (*124*). ASVs of the family *Rhizobiaceae* were classified as RH Nap$^+$, ASVs of the genus *Pseudomonas* were classified as PD Nar$^+$. Virtually no contamination (defined as ASVs classified as neither RH Nap$^+$ nor PD Nar$^+$) was observed ($< 0.2\%$ total counts, $< 0.8\%$ counts per condition).

Systematic measurement bias complicates the interpretation of 16S amplicon data as a reflection of biomass relative abundance (*125*). Therefore we sought to infer the relative abundances by biomass using a standard curve to infer conversion factors between ASV counts and biomass for each strain. Following Ref. 125, we assumed the biomass $x_*$ of each strain is proportional to the ASV counts $C_*$, with bias factors $b_*$:

$$\begin{aligned} x_{\mathrm{PD}} &= b_{\mathrm{PD}} C_{\mathrm{PD}} \\ x_{\mathrm{RH}} &= b_{\mathrm{RH}} C_{\mathrm{RH}} \end{aligned} \tag{6}$$

The relative abundance then becomes

$$f_{\mathrm{PD}} = \frac{C_{\mathrm{PD}}}{C_{\mathrm{PD}} + b_r C_{\mathrm{RH}}}. \tag{7}$$

where $b_r$ is the ratio of the bias factors $b_r = b_{\mathrm{RH}}/b_{\mathrm{PD}}$. Using measured counts for cultures with known PD Nar$^+$ relative abundances, we fit $b_r$ using the Levenberg-Marquardt algorithm (Fig. S5) and obtained a bias ratio of $4.3 \pm 0.7$. Eq. (7) was then used to infer biomass relative abundances from ASV counts. Error in inferred relative abundance comes from (a) error in the inference of $b_r$, and (b) variance between replicates. To account for (a), error in $b_r$ was inferred from the covariance of the fit, and corresponding uncertainty in relative abundance was calculated using the variance formula (*126*). To account for (b), standard deviation of relative abundance between biological replicates was calculated. The sum of the errors (a) and (b) was added in quadrature to compute the total error.

#### Measurement of mortality rates via plating

To measure and compare the effect of nitrite on mortality at pH $6.0$ and $7.3$, PD Nar$^+$ and RH Nap$^+$ were subjected to varying nitrite concentrations, and the number of viable cells over time was measured. This allowed us to infer a mortality rate as a function of pH and nitrite concentration.

First, plating media were selected to identify PD Nar$^+$ and RH Nap$^+$ separately. Preliminary experiments indicated that PD Nar$^+$ colonies were most clearly visible and well-defined on 1/10X TSB, $1.5\%$ agar plates ($1.5\,\mathrm{g/L}$ tryptone, $0.5\,\mathrm{g/L}$ soytone, $0.5\,\mathrm{g/L}$ sodium chloride, $15\,\mathrm{g/L}$ Bacto agar), and that RH Nap$^+$ colonies were most distinguishable on a defined medium with mannitol as the carbon source. Mannitol defined medium (MDM) was made using the same protocol as SDM, but $160\,\mathrm{mM}$ mannitol was substituted for succinate, and $1.5\%$ w/v agar was added.

Strains were then prepared for mortality experiments by preculturing as in phenotyping experiments (i.e., two stages of aerobic growth and one stage of anaerobic growth). After the anaerobic pre-culture step, $150\,\mu\mathrm{L}$ of culture of each strain normalized to $\mathrm{OD}_{600} = 0.04$ was passaged under anaerobic conditions to $1050\,\mu\mathrm{L}$ of SDM ($4\,\mathrm{mM}$ succinate, no nitrate) and incubated at $30\,^\circ\mathrm{C}$ and 950 RPM. Two pH conditions (6 and $7.3$) and four initial nitrite conditions ($0$, $0.438$, $0.875$, and $1.75\,\mathrm{mM}$) were tested.

Mortality rates were measured via time course of viable cell counts in each culture. $10\,\mu\mathrm{L}$ samples were taken from each condition at each time point. These samples were then serially tenfold diluted to achieve suspensions containing less than approximately $100$ cells; the number of dilutions necessary for each strain in each condition were determined by preliminary experiments. $10\,\mu\mathrm{L}$ of diluted samples were then pipetted onto solid medium, and streaked by tipping the plate at a roughly $45°$ angle until droplets approached the opposite edge of the plate. We eschewed mechanical streaking to avoid measurement bias. Plates were then covered and incubated at $30\,°\mathrm{C}$. Colonies in each streak were counted manually when distinct but not overgrown (approximately $1$ day for PD $\mathrm{Nar}^+$ on 1/10x TSB and $2$ days for RH $\mathrm{Nap}^+$ on MDM).

We mathematically combined cell density inferences from multiple streaks to reduce measurement error. Cell density inferred from each streak at dilution level $D_i$ can be written as

$$n_i = D_i C_i, \tag{8}$$

where $n_i$ is estimated cell density and $C_i$ is the number viable cells counted. To combine counts across dilution levels, weighted averages were used (*127*):

$$n_c = \frac{\sum_i \bar{n}_i w_i}{\sum_i w_i} \tag{9}$$

where $w_i = 1/\sigma_i^2$ are the weights and $\bar{n}_i$ and $\sigma_i$ are the average and standard deviation of the counts across biological replicates at dilution level $i$. The error on the combined counts is then given by

$$\sigma_c = \frac{1}{\sqrt{\sum_i w_i}}. \tag{10}$$

In the presence of growth, it is not possible to accurately measure mortality rate via plating. At pH $7.3$, growth was observed for all supplied nitrite values $I_0 > 0$, so so these conditions were discarded from analysis. At pH $6.0$, growth was observed at $I_0 = 0.5\,\mathrm{mM}$ (monotonic decrease in nitrite concentration, Fig. S6), so this condition was also discarded from further analysis. For the conditions showing no growth, mortality rate was inferred using a log-linear fit. By assuming the mortality rate is proportional to the number of cells, the number of viable cells can be expressed as follows:

$$n_c = n_{c,0} \exp\left(-r_d t\right) \tag{11}$$

where $r_d$ is the death rate, and $n_{c,0}$ is the number of viable cells at $t = 0$. Thus,

$$\ln(n_c) = \ln(n_{c,0}) - r_d t, \tag{12}$$

which is linear with respect to time. We, therefore, performed least-squares fits of $\ln(n_{c,0}) - r_d t$ to the logarithm of the measured number of viable counts and obtained $\ln(n_{c,0})$ and $r_d$ as fit parameters.

### Topsoil incubation experiments

#### Soil collection

To identify the taxa responsible for denitrification activity in natural soils using culture-free methods, topsoils were sampled across a range of pH values (5.0–7.1) from the Cook Agronomy Farm in September 2022 (Table S4). The Cook Agronomy Farm (CAF, 46.78°N, 117.09°W, 800 m above sea level) is a long-term agricultural research site located near Pullman, Washington, USA. CAF was established in 1998 as part of the Long-Term Agroecosystem Research (LTAR) network supported by the United States Department of Agriculture. Before being converted to an agricultural field, the site was zonal xeric grassland or steppe. CAF operates on a continuous dryland-crop rotation system comprising winter wheat and spring crops. CAF is located in the high rainfall zone of the Pacific northwest region and the soil type is classified as Mollisol (Naff, Thatuna and Palouse Series) (*128*).

Ten topsoils were collected from the eastern region of the CAF at a depth of 10–20 cm. Samples were collected within a diameter of 500 m to minimize the variation of edaphic factors other than pH. The large variation of soil pH comes from the long-term use of ammoniacal fertilizers and associated N transformations, combined with field-scale hydrologic processes that occur under continuous no-tillage superimposed over a landscape that has experienced long-term soil erosion. The pH measurements were made using a glass electrode in a 1:5 (soil to water) suspension of soil in Milli-Q filtered water. The ten soils had similar edaphic properties: 6–8% gravimetric water content (g/g), soil texture of silty clay or silty clay loam with 36–41% clay, and C:N ratio constnat at an approximate value of 12 with 1.1–1.8% total carbon (wt/wt) (Table S4).

#### Soil processing and incubation

To process soils for incubation, samples were sieved (<2 mm) to remove apparent plant roots and stones, and water content was measured (via drying at 105 °C for 24 h). To mimic autumn rainfall in the CAF area and stabilize microbial activity before beginning incubation, we rewetted the soil with sterile Milli-Q water at 40% water holding capacity for 2 weeks at room temperature. Soil slurries were then made by adding sterilized water to soil (2:1 w/w ratio of water to soil), which is close to natural state of soil saturated with water. Three replicates of each soil sample were then mixed with a concentrated sodium nitrate solution (nitrate$^{+}$ conditions) to yield an approximate 2 mM final concentration, and another three with sterile water (controls). These slurries were then transferred to 48-deepwell plates for incubation under anaerobic conditions (950 RPM, 30 °C) for approximately 96 h. Soil extracts were then prepared using a 2 M KCl solution, followed by 0.22 µm filtering to remove soil particles that may interfere with colorimetric assays. Colorimetric assays to measure the amount of nitrate consumed were carried out using a plate reader as described in earlier experiments. Post-incubation, endpoint samples were stored at −80 °C for subsequent DNA extraction and sequencing.

#### Sequencing and analysis of topsoil incubations

Genomic DNA was extracted from 400 µL incubation endpoint subsamples in a combined chemical and mechanical procedure using the PowerSoil DNeasy PowerSoil HTP 96 Kit (Qiagen, Hilden, Germany). Extraction was performed following the manufacturer's protocol, and extracted DNA was stored at −20 °C. To estimate the absolute abundance of bacterial 16S rRNA amplicons, known quantities of gDNA belonging to *Escherichia coli* B and *Parabacteroides* sp. TM425 (sample obtained from Duchossois Family Institute Commensal Isolate Library, Chicago, IL, USA) were added to the slurry subsamples before the DNA extraction step. DNA Library preparation was performed using the 16S Metagenomic Sequencing Library Preparation protocol with a 2-stage PCR workflow (Illumina, San Diego, CA, United States). The V3–V4 region was amplified using forward primer 341-b-S-17 (CCTACGGGNGGCWGCAG) and reverse primer 785-a-A-21 (GACTACHVGGGTATCTAATCC) (*129*). We confirmed using gel electrophoresis that the negative samples containing all reagents did not show visible bands after PCR amplification. Sequences were obtained on the Illumina MiSeq platform in a $2 \times 300$ bp paired-end run using the MiSeq Reagent Kit v3 (Illumina, San Diego, CA, United States). A standardized 10-strain gDNA mixture (MSA-1000, ATCC, Manassas, VA, USA) was sequenced as well to serve as a positive control.

Raw Illumina sequencing reads were stripped of primers, truncated at Phred quality score 2, trimmed to length 263 for forward reads and 189 for reverse reads (ensuring a 25-nucleotide overlap for most reads), and filtered to a maximum expected error of 4 based on Phred scores; this preprocessing was performed with USEARCH ver. 11.0 (*130*). The filtered reads were then processed with DADA2 ver. 1.26 following the developers' recommended pipeline (*123*). Briefly, forward and reverse reads were denoised separately, then merged and filtered for chimeras. For greater sensitivity, ASV inference was performed using the DADA2 pseudo-pooling mode, pooling samples by soil. After processing, the sequencing depth of denoised samples was $10^4$–$10^5$ reads per sample. Low-abundance ASVs were dropped, retaining 4466 ASVs for further analysis. (The samples used in this work were collected as part of a larger study; the exact abundance-filtering criterion was to retain ASVs with cumulative abundance of at least 1000 counts across the 902 samples of this larger dataset.) Taxonomy was assigned by DADA2 using the SILVA database ver. 138.1, typically at genus level, but with species-level attribution recorded in cases of a 100% sequence match.

As an internal control, we verified that the ASVs corresponding to the two spiked-in genera *Escherichia-Shigella* and *Parabacteroides* were highly correlated with each other as expected ($c = 0.94$). These ASVs were removed from the table and combined into a single reference vector of "spike-in counts". The spike-in counts constituted $5.5 \pm 2.5\%$ of total reads in each sample.

For subsequent analysis, the raw ASV counts were augmented by a pseudocount of 0.5 and divided by the per-sample spike-in counts, yielding values that can be interpreted as the absolute biomass of each taxon (up to a factor corresponding to the copy number of the 16S operon), measured in units where 1 means as many 16S fragments as the number of DNA molecules in the spike-in.

##### Identifying and genotyping ASVs enriched on nitrate in topsoil incubations

To identify the ASVs enriched in nitrate treatments versus the no-nitrate controls, it was necessary to determine what change in recorded abundance constitutes a significant change, relative to what might be expected for purely stochastic reasons. The relevant null model would combine sampling and sequencing noise with the stochasticity of ecological dynamics over a 4-day incubation, and cannot be derived from first principles. However, since all measurements were performed in triplicate with independent incubations, the relevant null model can be determined empirically. The deviations of replicate-replicate comparisons from 1:1 line were well-described by an effective model combining two independent contributions, a Gaussian noise of fractional magnitude $c_{\text{frac}}$ and a constant Gaussian noise of magnitude $c_0$ reads, such that repeated measurements (over biological replicates) of an ASV with mean abundance $n$ counts are approximately Gaussian-distributed with a standard deviation of $\sigma(c_0, c_{\text{frac}}) = \sqrt{(c_{\text{frac}}n)^2 + c_0^2}$ counts (Fig. S11). In this expression, $c_{\text{frac}}$ was estimated from moderate-abundance ASVs ($> 50$ counts) for which the other noise term is negligible; and $c_0$ was then determined as the value for which 67% of replicate-replicate comparisons are within $\pm\sigma(c_0, c_{\text{frac}})$ of each other, as expected for 1-sigma deviations. This noise model was inferred separately for each soil, as the corresponding samples were processed independently in different sequencing runs; the parameters across 10 soils were $c_{\text{frac}} = 0.22 \pm 0.03$ and $c_0 = 11 \pm 3$ counts. The model was used to compute the z-scores for the enrichments of absolute ASV abundances in nitrate treatments against no-nitrate controls (three independent z-scores, from triplicate treatments). Significantly enriched ASVs were identified in each sample as those with z-scores greater than $z = \Phi^{-1}(1 - \alpha/2/n_{ASV})$, where $\Phi^{-1}(x)$ is the inverse CDF of the standard normal distribution, $\alpha = 0.01$, and $n_{ASV}$ as the number of nonzero ASVs in a given sample. This critical z-score ($z = 4.51 \pm 0.04$) corresponds to a two-tailed Bonferroni-corrected hypothesis test at significance level $\alpha$ under the null hypothesis that counts in the nitrate$^{+}$ and control conditions are drawn from the same distribution. These analyses were performed using custom MATLAB scripts (Mathworks, Inc), which are available on the GitHub data repository for the present manuscript; for additional technical details, the reader is referred to the detailed comments in these scripts.

In order to determine which significantly enriched ASVs are likely to represent denitrifiers, PICRUSt2 ver. 2.5.2 (*50*) was used to assign putative genotypes. PICRUSt2 matches input 16S rRNA sequences to genotypes using a curated reference genome database. A list of inferred KEGG orthologs for each ASV was computed using the default parameters of the PICRUSt2 pipeline. An ASV was classified as a denitrifier if KEGG orthologs for any denitrification reductase (*narG*/K00370, *napA*/K02567, *nirS*/K15864, *nirK*/K00368, *norB*/K04561, *nosZ*/K00376) were inferred *and* if the DNRA nitrite reductase *nrfA*/K03385 was not inferred. ASVs were classified as performing DNRA if *nrfA*/K03385 was inferred, and as neither performing denitrification nor DNRA otherwise.

### Supplementary Information

### 1 Consumer resource model

To fit metabolite dynamics data to a consumer-resource model (CRM), we used the protocol described by Gowda *et al.* (*23*). Briefly, the CRM is

$$\begin{aligned} \frac{dx}{dt} &= \left(\gamma_A r_A \frac{A}{K_A + A} + \gamma r_I \frac{I}{K_I + I}\right) x \\ \frac{dA}{dt} &= -\sum_{i=1}^{N} r_A \frac{A}{K_A + A} x \\ \frac{dI}{dt} &= \sum_{i=1}^{N} \left(r_A \frac{A}{K_A + A} - r_I \frac{I}{K_I + I}\right) x. \end{aligned} \tag{13}$$

where $x$ is biomass of a single strain, $A$ and $I$ are the concentrations of nitrate and nitrite respectively, $\gamma_A$ and $\gamma_I$ are biomass yields, $r_A$ and $r_I$ are per capita reduction rates, and $K_A$ and $K_I$ are affinities (*23*), on nitrate and nitrite respectively. As motivated by Ref. 23, the affinity parameters were set to $0.01\,\mathrm{mM}$ because the true values are likely on the order of µM and therefore cannot substantially affect dynamics at $\mathrm{mM}$ concentrations.

#### 1.1 pH 7.3 fits and co-culture predictions

We used the CRM (Eq. 13) to show that the results of co-culture competition experiments at at pH 7.3 are consistent with the environmental filtering hypothesis. To do this, we used the CRM to fit monoculture data and then predict the outcome of a co-culture competition experiment. RH $\mathrm{Nap}^+$, the strain enriched at pH 7.3, is expected to dominate in co-culture at pH 7.3 based on monoculture growth dynamics, suggesting that it is better *individually* adapted to a pH 7.3 environment.

PD $\mathrm{Nar}^+$ and RH $\mathrm{Nap}^+$ were grown in monoculture for initial nitrate and nitrite concentrations $(A_0, I_0) = (2,0), (1,0), (0,2), (0,1)$, and yield inferred following Ref. 23. With these yield values fixed ($\gamma_{A,\mathrm{Nar}^+} = 0.018\,\mathrm{OD/mM}$, $\gamma_{I,\mathrm{Nar}^+} = 0.022\,\mathrm{OD/mM}$, $\gamma_{A,\mathrm{Nap}^+} = 0.019\,\mathrm{OD/mM}$, and $\gamma_{I,\mathrm{Nap}^+} = 0.029\,\mathrm{OD/mM}$) and affinity parameters set to low values, we then numerically integrated the CRM to perform a least-squares fit of the remaining parameters $r_A$ and $r_I$ to the dynamics of nitrate and nitrite ($r_{A,\text{Nar}^+} = 5.1\,\text{mM/OD/h}$, $r_{I,\text{Nar}^+} = 2.8\,\text{mM/OD/h}$, $r_{A,\text{Nap}^+} = 2.6\,\text{mM/OD/h}$, and $r_{I,\text{Nap}^+} = 7.1\,\text{mM/OD/h}$). These fits are shown in Fig. S7A-H.

To predict co-culture relative abundances, we used the following extended CRM,

$$\begin{aligned} \frac{dx_i}{dt} &= \left(\gamma_A^i r_A^i \frac{A}{K_A + A} + \gamma_I^i r_I^i \frac{I}{K_I + I}\right) x_i \quad \text{for } i = \text{Nar}^+, \text{Nap}^+ \\ \frac{dA}{dt} &= -\sum_{i=1}^{N} r_A^i \frac{A}{K_A + A} x_i \\ \frac{dI}{dt} &= \sum_{i=1}^{N} \left(r_A^i \frac{A}{K_A + A} - r_I^i \frac{I}{K_I + I}\right) x_i, \end{aligned} \tag{14}$$

which simulates a co-culture by modeling the biomass $x_i$ of each strain $i$, and summing the contributions of each strain to the consumption/production of nitrate and nitrite. We integrated this model for each cycle using the parameters fit to monoculture growth dynamics, as described in Ref. 23. Serial dilutions with $72\,\text{h}$ cycles, eight-fold dilutions, and $2\,\text{mM}$ of initially supplied nitrate were simulated. We performed this simulation for a total of $4$ cycles over a range of initial relative abundances. Relative abundances ($x_i/\sum_i x_i$) were computed for each phenotype at the end of each cycle. In all simulated enrichments $K_A = K_I = 0.01\,\text{mM}$. The results of these simulations are shown Fig. S7I. For all initial relative abundance conditions, the model predicts that RH Nap$^+$ dominates at pH $7.3$ based on the monoculture phenotypes of both strains. This is consistent with the environmental filtering hypothesis.

#### 1.2 pH $6.0$ fits and monoculture predictions

We also used a consumer resource model to illustrate the effect of nitrite toxicity on growth dynamics at pH $6.0$.

In order to determine yields on nitrate and nitrite using bulk measurements, it is typically informative to grow strains with varying initial nitrite concentrations (*23*). However, since high concentrations of nitrite are toxic at low pH, it is difficult to use this approach to measure yields at pH $6.0$ (Fig. 4). We, therefore, measured yields at pH $7.3$ and made the assumption that these values apply to pH $6.0$ as well. PD Nar$^+$ was grown for initial nitrate and nitrite concentrations

$(A_0, I_0) = (2, 0), (1, 0), (0, 2), (0, 1)$ at pH 7.3, and yield inferred following Ref. 23. These estimates ($\gamma_A = 0.018\,\mathrm{OD/mM}$ and $\gamma_I = 0.022\,\mathrm{OD/mM}$) predicted well (6% error) the endpoint OD for the $(A_0, I_0) = (1, 0)$ condition at pH $6.0$, for which we expect the effect of nitrite toxicity is negligible. With these yield values fixed and affinity parameters set to low values, we then numerically integrated the CRM to perform a least-squares fit of the remaining parameters $r_A$ and $r_I$ to the dynamics of nitrate and nitrite in the $(A_0, I_0) = (1, 0)$ pH $6.0$ condition using nonlinear optimization. These best fit parameters ($r_A = 5.8\,\mathrm{mM/OD/h}$ and $r_I = 1.6\,\mathrm{mM/OD/h}$) were saved and used to predict the metabolite dynamics of the pH $6.0$ $(2, 0)$ condition via numerical integration.

### 2 Statistical analysis of reference denitrifier genomes

We studied patterns of gene presence-absence across a diverse collection of denitrifier genomes, in particular in order to investigate whether the nitrate reductase genes *narG* and *napA* are typically found together or independently.

First we curated a library of prokaryotic denitrifying genomes from the KEGG reference genome database (*131–133*), selecting genomes containing at least one denitrification reductase (*narG*, *napA*, *nirS*, *nirK*, *cNor*, *qNor*, *nosZ*) and not containing genes for dissimilatory nitrate reduction to ammonia (*nrfA*, *nrfH*). Applying this filter to the database resulted in $2154$ genomes, and for each we recorded the presence or absence of each of the $7$ denitrification reductases in a binary matrix. We note that this analysis, in contrast to the earlier analysis of the global topsoil microbiome, distinguishes between the nitric oxide reductases *cNor* and *qNor*. *qNor* is a fusion of the subunits *norB* and norC that make up *cNor* (*134*). It is routine to separately identify and annotate *cNor* and *qNor* in whole genome assemblies. However doing the same in the global topsoil microbiome was not straightforward, because the low depth of sequencing in this dataset prevented a reliable assembly of sufficiently long sequences to distinguish *cNor* and *qNor*.

We took a phylogenetically-weighted adjusted mutual information approach to quantifying the relationships between genes in genomes. First, for two binary vectors $\mathbf{x}$, $\mathbf{y}$, here representing the presence/absence of two genes across the collection of genomes, mutual information (MI) is defined by

$$\text{MI}(\mathbf{x}, \mathbf{y}) = \sum_{i} \sum_{s \in \{0,1\}} \sum_{t \in \{0,1\}} p(x_i = s, y_i = t) \log \left( \frac{p(x_i = s, y_i = t)}{p(x_i = s) p(y_i = t)} \right) \tag{15}$$

MI indicates the amount of information about $\mathbf{x}$ contained in $\mathbf{y}$, and vice versa; MI is largest when $\mathbf{x} = \mathbf{y}$, and MI $\approx 0$ when the underlying distributions of entries in $\mathbf{x}$ and $\mathbf{y}$ are independent. MI is bounded above by the marginal entropy: $MI(\mathbf{x}, \mathbf{y}) \leq \min(H(\mathbf{x}), H(\mathbf{y}))$, where entropy $H(\mathbf{x}) = -\sum_i \sum_{s \in \{0,1\}} p(x_i = s) \log p(x_i = s)$.

Because the vectors $\mathbf{x}$ and $\mathbf{y}$ are finite, MI is likely to take a non-zero value simply due to chance. The adjusted mutual information (AMI) corrects this effect by offsetting MI by the expected MI under the hypothesis of independence:

$$\text{AMI}(\mathbf{x}, \mathbf{y}) = \frac{\text{MI}(\mathbf{x}, \mathbf{y}) - E\{\text{MI}(\mathbf{x}, \mathbf{y})\}}{\min(H(\mathbf{x}), H(\mathbf{y})) - E\{\text{MI}(\mathbf{x}, \mathbf{y})\}}. \tag{16}$$

The result is a measure that takes the value $1$ if $\mathbf{x} = \mathbf{y}$ and $0$ if the $\text{MI}(\mathbf{x}, \mathbf{y})$ is equal to the MI due to chance alone. The expected MI was estimated from pairs of randomly-permuted versions of the vectors $\mathbf{x}$ and $\mathbf{y}$ (*135*). Finally, in order to differentiate positively and negatively correlated relationships between genes, we assigned a sign (+/-) to the value of AMI depending on the sign of the Pearson correlation between $\mathbf{x}$ and $\mathbf{y}$.

One crucial consideration is that the selection of reference genomes is phylogenetically biased. Some species are more extensively represented in the KEGG reference genome database (e.g., strains of *E. coli*) than others. To account for this bias, we calculated phylogenetic weights for each genome by constructing a phylogenetic tree using 16S rRNA sequences from each genome (annotated by KO K01977). The weight $w_i$ for genome $i$ was defined as the inverse of genome $i$'s phylogenetic cluster size, i.e., the number genomes with 16S rRNA sequence similarities larger than $1 - \delta$ (*136*), where $\delta$ is the fraction of nucleotides that differ between a pair of 16S rRNA genes. Intuitively, this approach assigns smaller weights to genomes in densely sampled branches of the phylogenetic tree to reduce this branch's contribution, and larger weights to genomes in less well-sampled branches. We applied these weights to all sums over genomes in order to compute phylogeny-weighted AMI (Fig. S9). Here we chose $\delta = 0.1$, and observed that the results do not depend strongly on the value of this parameter.

Fig. S9 shows that the genes *narG* and *napA* have a negative phylogeny-weighted AMI, meaning that genomes possessing *narG* tend not to possess *napA*, and vice versa.

### 3 Simulations of enrichment experiments with trade-offs and toxicity

We used simulations of a CRM to investigate the role of physiological trade-offs and toxicity in community assembly across a pH gradient. These simulations support our Discussion of the potential role of trade-offs in the observed patterns.

#### 3.1 Model

We extended the CRM described above (Eq. 14) to $N$ strains:

$$\begin{aligned}
\frac{dx_i}{dt} &= \left(\gamma_A^i r_A^i \frac{A}{K_A + A} + \gamma_I^i r_I^i \frac{I}{K_I + I} - r_d^i(I)\right) x_i \quad \text{for } i = 1, \ldots, N \\
\frac{dA}{dt} &= -\sum_{i=1}^{N} r_A^i \frac{A}{K_A + A} x_i \\
\frac{dI}{dt} &= \sum_{i=1}^{N} \left(r_A^i \frac{A}{K_A + A} - r_I^i \frac{I}{K_I + I}\right) x_i.
\end{aligned} \tag{17}$$

and added a nitrite concentration-dependent death rate,

$$r_d^i(I) = r_{\text{d,max}}^i \frac{1}{1 + \exp^{-\alpha_d^i (I - I_{1/2}^i)}} \tag{18}$$

where $I_{1/2}$ sets the nitrite concentration $I$ at which the death rate is half of its maximum value ($r_{\text{d,max}}$), and $\alpha_d$ controls the rate at which the death rate increases. Thus this death rate term takes values between 0 (when $I \ll I_{1/2}$) and $r_{\text{d,max}}$ (when $I \gg I_{1/2}$). For these simulations, we used $\alpha_d^i = 10^4\,\text{mM}^{-1}$. In this limit, the death rate is well-approximated by a Heaviside step function,

$$r_d^i(I) \approx r_{\text{d,max}}^i \mathcal{H}(I_{1/2}^i - I). \tag{19}$$

#### 3.2 Phenotypic constraints

Modelled phenotypes were constrained based on (a) a tradeoff between growth rate on nitrate and nitrite, and (b) a tradeoff between growth rate on nitrate and a sensitivity to nitrite toxicity (Fig. S12A).

A tradeoff between nitrate and nitrite growth is both well-supported in the literature (*33*) and consistent with our experimental results. In particular, we observed that PD $Nar^+$ grows more quickly on nitrate than nitrite, whereas RH $Nap^+$ grows more quickly on nitrite than nitrate (PD $Nar^+$ accumulates nitrite whereas RH $Nap^+$ does not, Figs. 3, S7). To instantiate this tradeoff mathematically, we ensured that

$$0 = d - b(r_A/a)^k - c(r_I/a)^k \tag{20}$$

where $r_A$ and $r_I$ are the CRM per capita uptake rates of nitrate and nitrite, respectively, $d$ and $a$ set the overall scaling, $b$ and $c$ control symmetry, and $k$ controls the curvature of the tradeoff (Fig. S12A). For these simulations, $b = c = 1$ (symmetric tradeoff). $d = 1$ and $a = 5$ to approximate measured rates.

The second constraint imposed on our system, a tradeoff between nitrate growth rate and nitrite-concentration dependent death rate is empirically motivated. We observed that the strain that grows preferentially on nitrate, PD $Nar^+$, experiences mortality when exposed to high levels of nitrite toxicity. This is not the case for the strain that grows preferentially on nitrite, RH $Nap^+$ (Fig. 4B). To instantiate this tradeoff mathematically, we included the death term $r_d(I)$ for strains with $r_A$ greater than some threshold $r_{A,\text{thresh}}$ (dashed line, Fig. S12A). For the simulations in Fig. S12, $r_{A,\text{thresh}} = 2\,\text{mM/OD/h}$.

#### 3.3 Simulated enrichment

To use the model to simulate possible community outcomes, we performed an *in silico* enrichment experiment, analogous to the one described *in vitro*. We numerically integrated Eq. 17 for a community initially composed of 20 constrained phenotypes ($r_A$, $r_I$) (Eq. 20). Serial dilutions with 72 h cycles, eight-fold dilutions, and $2\,\text{mM}$ of initially supplied nitrate were simulated. We performed this simulation for a total of $12$ cycles (and $4$ cycles for co-culture enrichments in Fig. S12E & F). Relative abundances $x_i/\sum_i x_i$ were computed for each phenotype at the end of each cycle. In all simulated enrichments $K_A = K_I = 0.01\,\text{mM}$ and $\gamma_A = \gamma_I = 0.02\,\text{OD}/\text{mM}$.

#### 3.4 Parameter space scan

Enrichment was simulated for many values of the curvature parameter $k$ (Eq. 20) and toxicity threshold parameter $I_{1/2}$ (Eq. 18). Increasing $k$ corresponds to an increase in the leniency of the physiological tradeoff (i.e., increase in tradeoff curvature, Fig. S12A), while increasing $I_{1/2}$ decreases the extent of nitrite toxicity. $I_{1/2}$, therefore, functions as a proxy for pH. The region of parameter space relevant to the experimental system is shown in Fig. S12B, and a more complete scan is shown in Fig. S13. This scan demonstrates that the results shown in Fig. S12B are not sensitive to the choice of $r_{d,\text{max}}$ or $r_{A,\text{thresh}}$, and are therefore robust to the choice of parameter.

#### 3.5 Classification of endpoint community structure

Fig. S12B and Fig. S13 show the classification (colors) of the simulated endpoint community structure by relative abundance distribution. We classified communities by the locations of the peaks in the relative abundance distributions (Fig. S12C, D), and distinguished between specialist phenotypes, for which either $r_A = 0$ or $r_I = 0$, and generalist phenotypes, for which $r_A > 0$ and $r_I > 0$. Thus, if the relative abundance distribution contains two peaks for which $r_A > 0$ and $r_I > 0$, we referred to it as a two-generalist regime, and if the peaks are at $r_A > 0$ and $r_I = 0$, we referred to it a one-generalist, one-specialist regime, and so on. If there is only one specialist, we distinguished between nitrate specialists ($r_I = 0$) and nitrite specialists ($r_A = 0$).

### Supplementary Figures

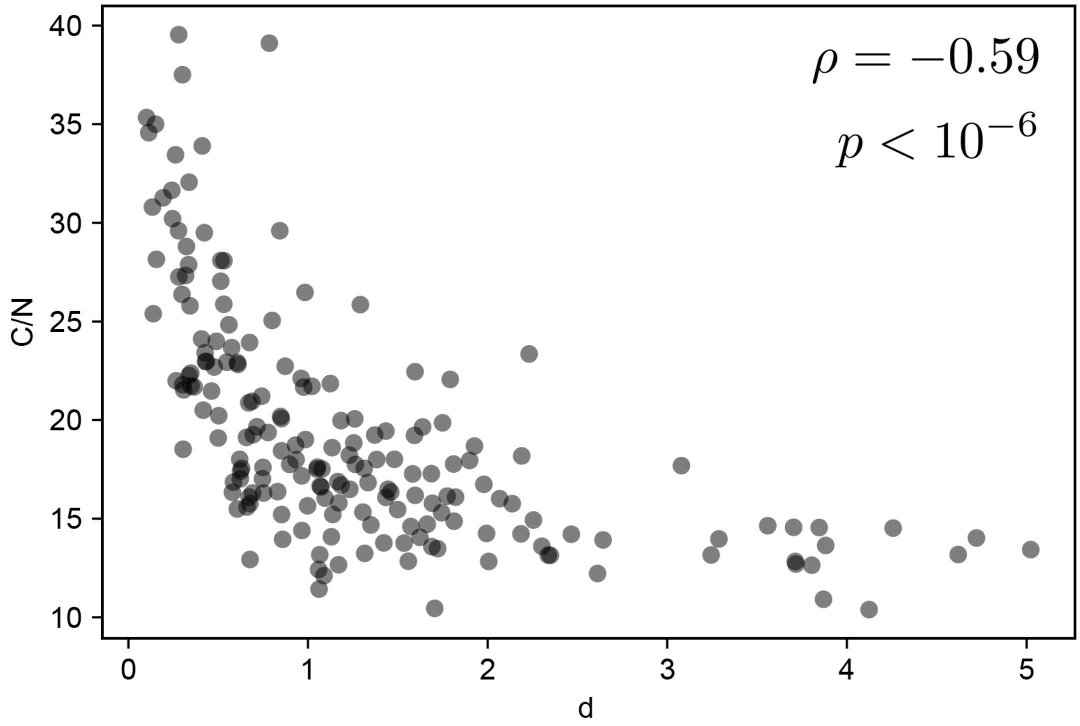


Figure S1: **C/N ratio correlates negatively with denitrification pathway magnitude.** Unit-invariant singular value decomposition (uiSVD) was used to decompose denitrification reductase (*narG*, *napA*, *nirS*, *nirK*, *norB*, *nosZ*) relative abundances from the global topsoil microbiome into contributions due to pathway magnitude and composition (Fig. 1A-D). Pathway magnitude ($d$) most strongly correlated with C/N ratio ($\rho = -0.59$, $p < 10^{-6}$ via one-tailed randomization test; Fig. 1E).

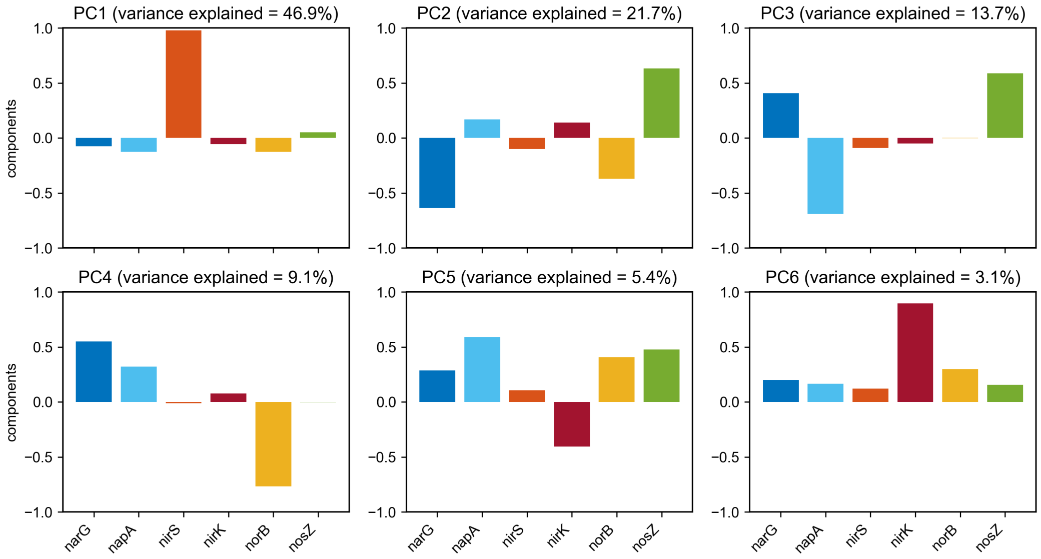


Figure S2: **Principal components resulting from the uiSVD decomposition of the global topsoil microbiome.** Bars show the loadings of each gene in the six principal components (PCs) resulting from the uiSVD decomposition of denitrification reductase (*narG*, *napA*, *nirS*, *nirK*, *norB*, *nosZ*) relative abundances from the global topsoil microbiome.

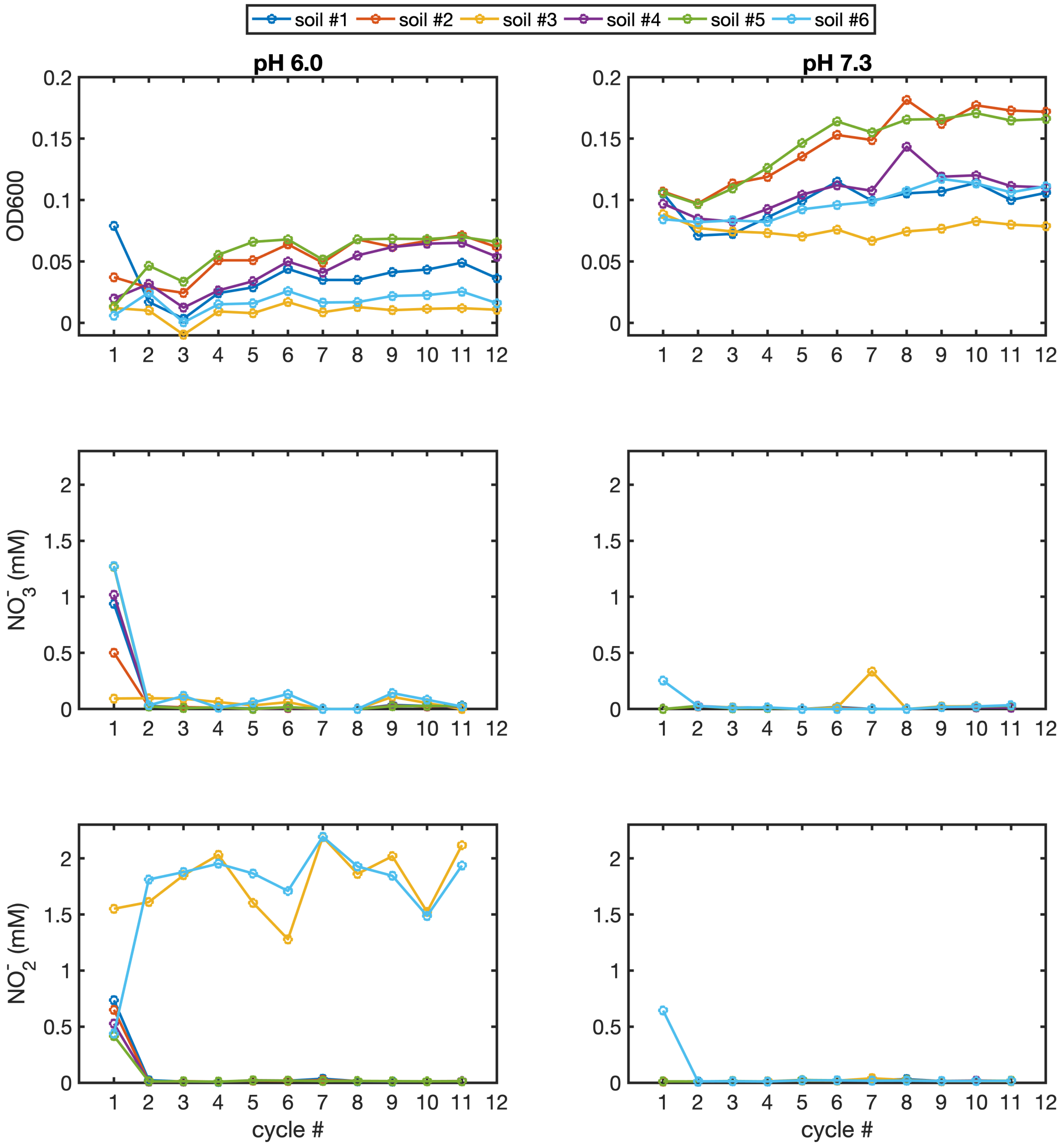


Figure S3: **Abundance and denitrification dynamics in primary soil enrichment experiment.** Enrichment cultures were performed by serially passaging communities extracted from six soil samples in defined medium buffered at two pH conditions, pH 6.0 and 7.3 for a total of 12 cycles. Optical densities, nitrate, and nitrite concentrations are shown at the endpoint of each cycle. Cultures initiated with soils #3 and 6 at pH 6.0 accumulated significant concentrations of nitrite at the end of each cycle.

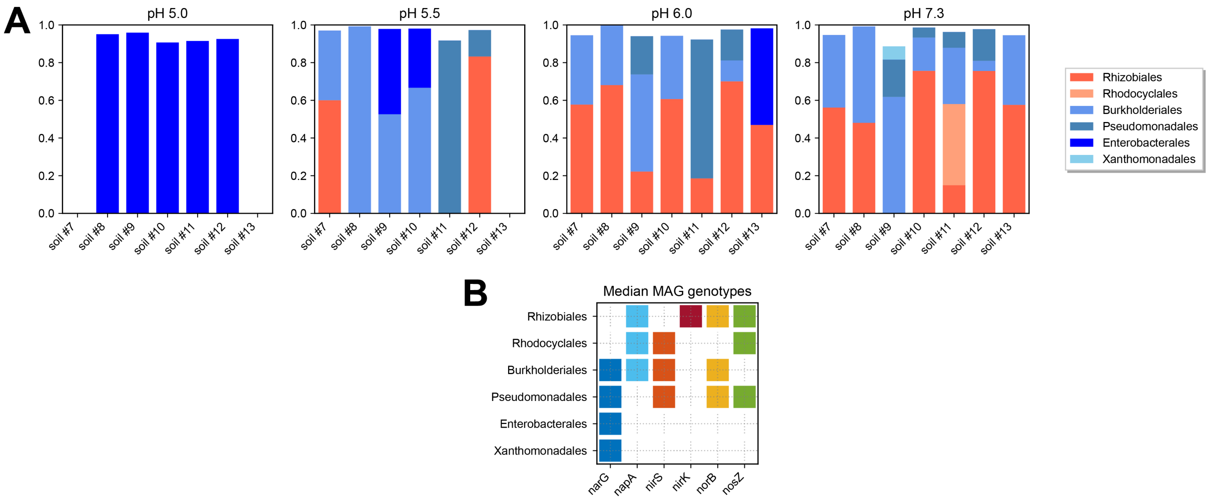


Figure S4: **Enrichment across a broader range of pH values.** Additional enrichments were performed at pH 5.0, 5.5, 6.0, and 7.3 and the endpoint cultures were shotgun sequenced to infer taxonomic composition and genotypes. **(A)** Endpoint community compositions of the enrichments inferred via 16S miTAGs are shown. Taxa with $Nar^+$ genotypes are indicated in shades of blue, while taxa that possess Nap and not Nar are indicated in shades of red. Compositions are shown at the level of taxonomic order, and taxa present at a level of less than $1\,\%$ are omitted. **(B)** Median denitrification reductase genotypes inferred via annotation of metagenome assembled genomes are shown.

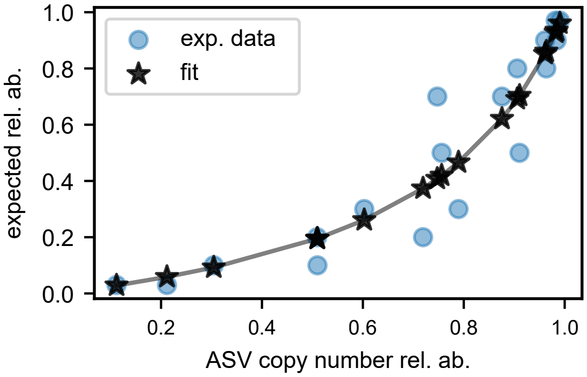


Figure S5: **Fit of bias ratio to infer relative abundances from ASV counts.** Blue dots show controls for which relative abundance is known (vertical axis) and ASV copy number relative abundance was measured (horizontal axis), with $n = 2$ technical replicates at each expected relative abundance. Black stars indicate best fit values of relative abundances inferred from ASV copy number relative abundance via a fit of Eq. 7 to the data ($b_r = 4.3 \pm 0.7$, where the error is the standard deviation of the fit value).

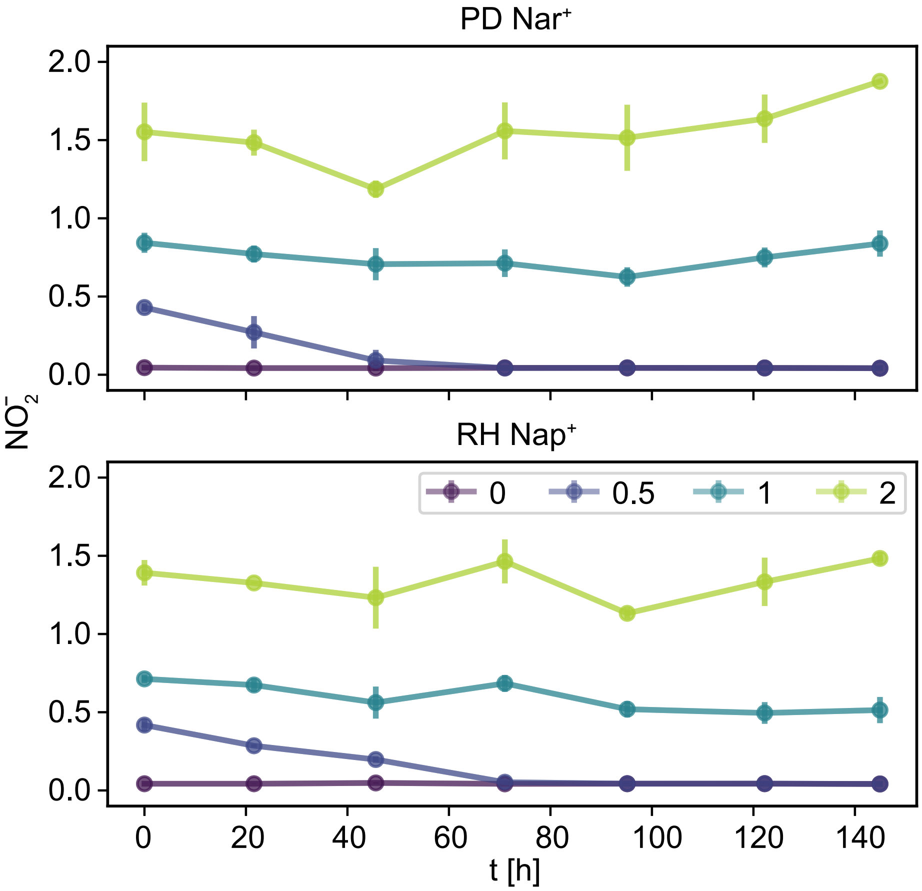


Figure S6: **Metabolite dynamics during mortality rate measurement.** Metabolite dynamics corresponding to the mortality rate experiment (main text Fig. 4B) are shown. All curves started with $0\,\mathrm{mM}$ $NO_3^-$, and the colors indicate the initial $\mathrm{mM}$ concentration of $NO_2^-$. For both strains, $NO_2^-$ was only reduced in the $0.5\,\mathrm{mM}$ initial condition. At higher $NO_2^-$ concentrations toxicity inhibits metabolic activity in both strains. Points and error bars show means and standard deviations over technical replicates ($n = 3$) in each condition.

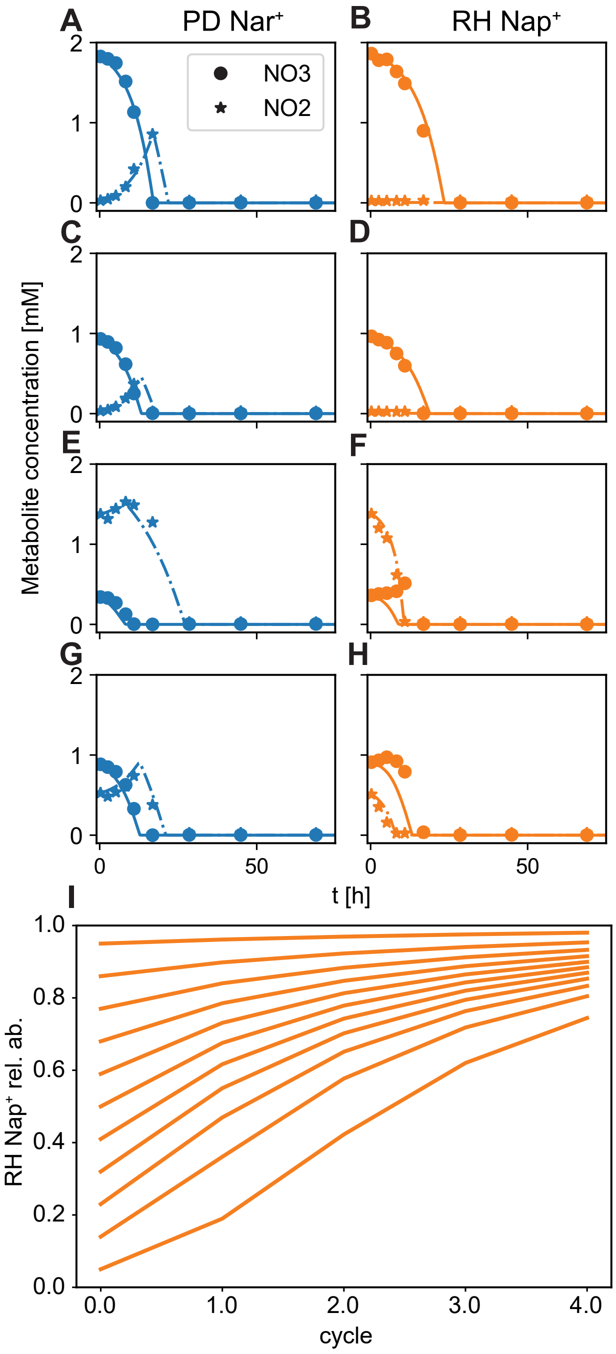


Figure S7: **pH 7.3 monoculture data is consistent with environmental filtering. (A)-(H)** Consumer resource model (Eq. 13) fit to monoculture metabolite data at pH 7.3. Dots indicate nitrate concentrations, stars indicate nitrite concentrations, solid lines show fits to nitrate dynamics, and dash-dot lines show fits to nitrite dynamics. All concentrations are averaged over technical replicates ($n = 3$). Panels (A, C, E, G) are fits to PD Nar$^+$ data, while panels (B, D, F, H) are fits to RH Nap$^+$ data. To infer nitrate and nitrite reduction rates independently fits were performed for a number of different initial conditions ($[NO_3^-]$, $[NO_2^-]$). Panels (A) and (B) correspond to (1.75, 0), (C) and (D) to (0.875, 0), (E) and (F) to (0.4375, 1.3125), (G) and (H) to (0.875, 0.4375). All concentrations are reported in units of mM. **(I)** Co-culture relative abundance prediction based on monoculture phenotypes. RH Nap$^+$ is predicted to approach a relative abundance of 1 for all initial conditions at pH 7.3. This is consistent with the environmental filtering hypothesis and is observed experimentally (main text Fig. 5F)

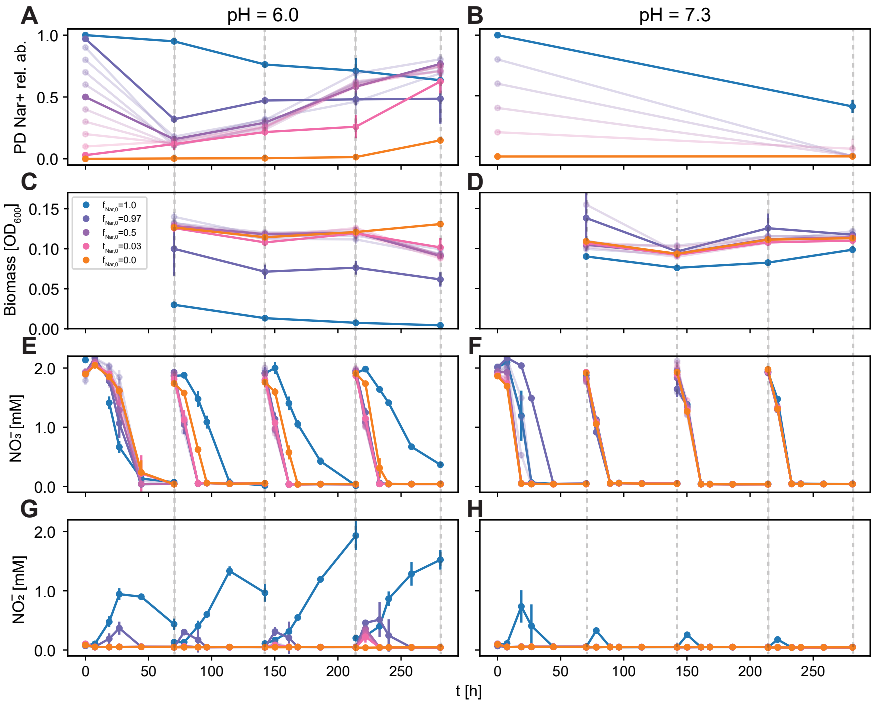


Figure S8: **Co-culture enrichment biomass and metabolite dynamics.** Details of the PD Nar$^+$ and RH Nap$^+$ co-culture experiment shown in main text Fig. 5. (**A**) PD Nar$^+$ relative abundance dynamics at pH 6.0 are shown. $f_{0,Nar^+} = 0, 0.03, 0.5, 0.97$ and 1 are highlighted. Due to small levels of cross-contamination between pure and mixed cultures, $f_{Nar^+}$ increases from 0 and decreases from 1. Although this was unintentional, it indicates that each of these strains is invasible by the other in this condition, providing more evidence that they coexist. (**B**) PD Nar$^+$ relative abundance dynamics at pH 7.3 are shown. (**C**) Endpoint biomass dynamics, measured via absorbance at 600 nm, are shown for each cycle at pH 6.0. The $f_{0,Nar^+} = 0$ condition produces much less biomass than the other conditions, as expected. (**D**) Endpoint biomass dynamics are shown for each cycle at pH 7.3. (**E, G**) $NO_3^-$ and $NO_2^-$ dynamics are measured using a Griess assay (*137*) and shown at pH 6.0. Aside from the $f_{0,Nar^+} = 0$ condition, for which biomass is very low (C), increasing $f_{Nar^+}$ (A) corresponds to increasing nitrite accumulation. (**F, H**) Metabolite dynamics are shown at pH 7.3. Decreasing $f_{Nar^+}$ (B) corresponds to decreasing nitrite accumulation. Points and error bars show means and standard deviations over technical replicates ($n = 3$ or $n = 4$) in each condition (see available code and data for detail).

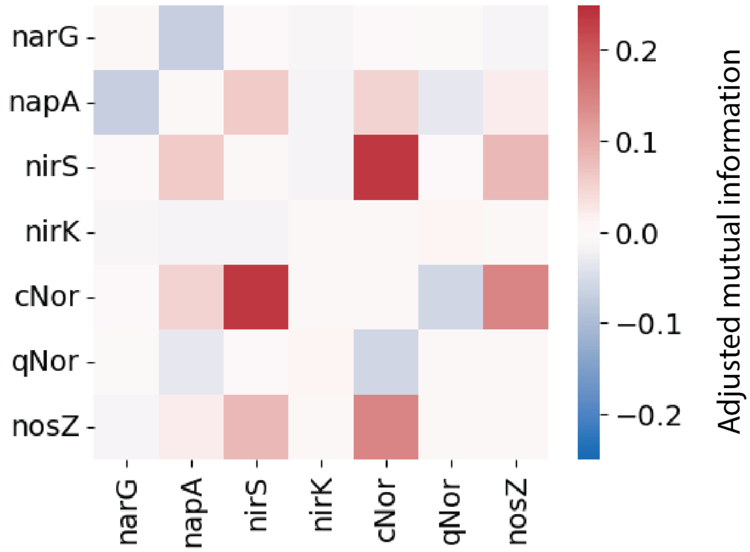


Figure S9: **Co-evolution of denitrification genes.** Phylogeny weighted adjusted mutual information (AMI) of each pair of main denitrification genes. The two nitrate reductases, *narG* and *napA* are negatively correlated. $\delta = 0.1$.

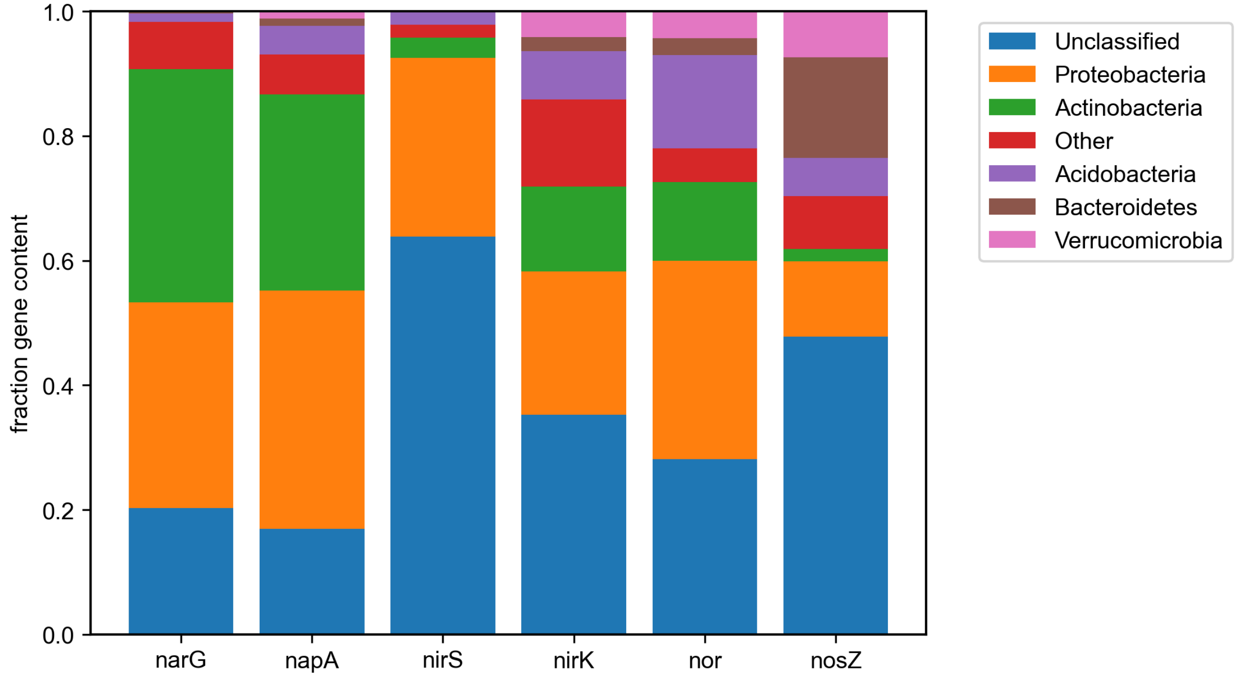


Figure S10: **Taxonomic classification of denitrification reductases in global topsoil microbiome.** Reads annotated as denitrification reductases (*narG*, *napA*, *nirS*, *nirK*, *norB*, and *nosZ*) in metagenomic sequencing of the global topsoil microbiome were combined and then taxonomically classified using the Kaiju web server (*104*). The fraction of reads are shown for phyla present at a level of 5% or higher for any reductase, with the remainder plotted as "other" (red). Classifiable reads were assigned to Proteobacteria (orange) more often than any other phylum. Proteobacterial reads comprise 12%–38% of total reads depending on the reductase.

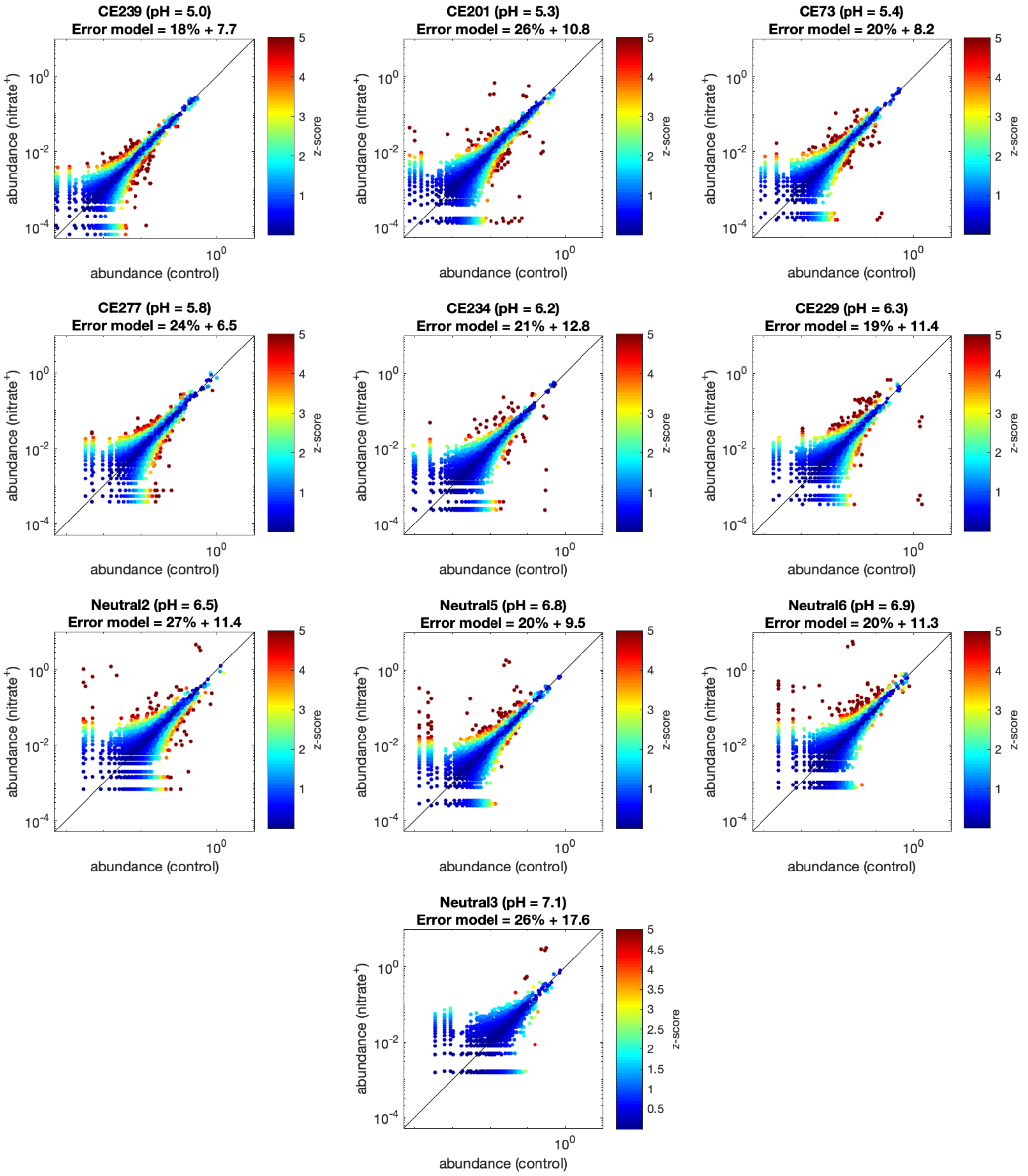


Figure S11: **Differential abundance analysis of ASVs in topsoil microcosms.** ASV abundances in controls versus nitrate$^+$ conditions are shown for each soil sample, with the diagonal line indicating equal abundance values. Colors indicate the magnitude of z-score, a measure of differential abundance between conditions. Parameters of an error model ($c_{\text{frac}}$ and $c_0$, respectively) inferred for each soil condition are given. Abundance/z-score values for $N = 3$ replicates of each soil sample are plotted.

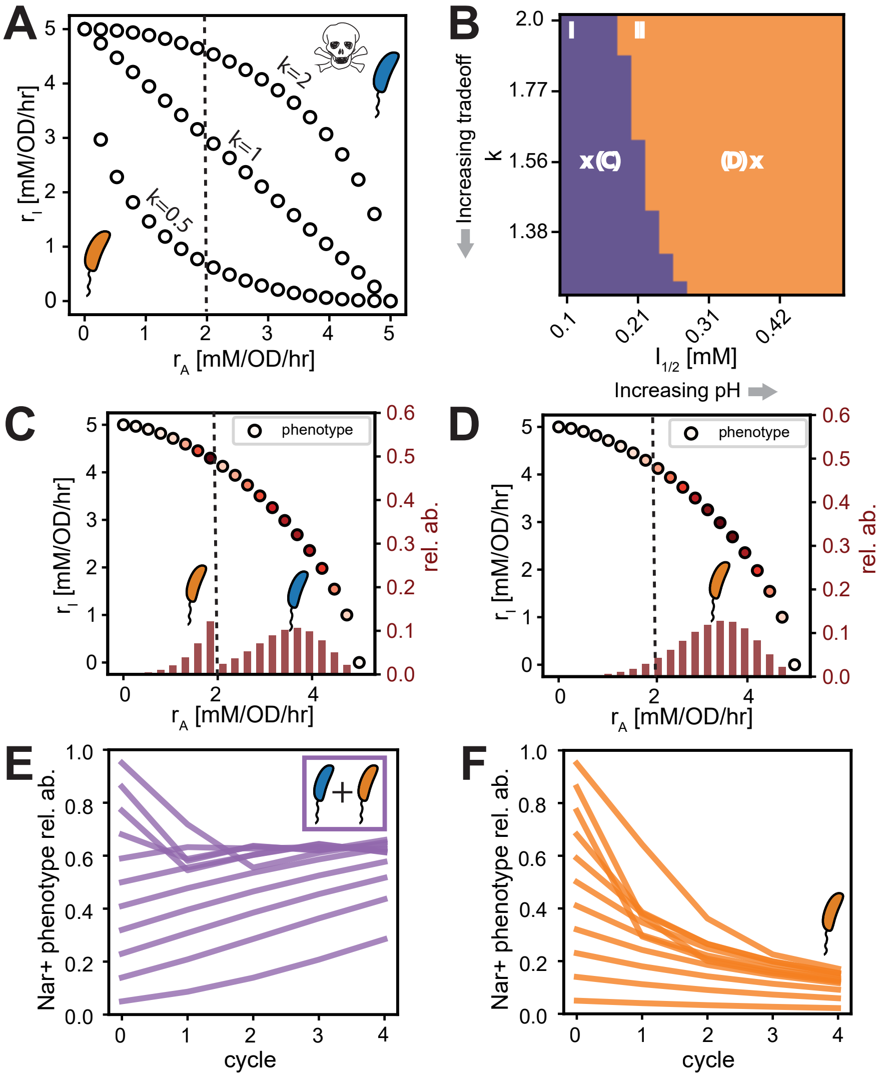


Figure S12: See following page.

Figure S12: **Simulations of enrichments of diverse communities with phenotypes subjected to metabolic tradeoffs and toxicity.** (**A**) The model is illustrated. The trade-off in nitrate reduction rate $r_A$ and nitrate reduction rate $r_I$ is shown (left panel). Trade-off curvature is controlled by the parameter $k$ (Supplementary Information). A point on this Pareto front indicates the phenotype of one strain ($r_A$, $r_I$). A community of Pareto optimal strains is illustrated by the open circles. In addition to the trade-off, fast nitrate reducers (strains to the right of the dashed line), are killed by nitrite concentrations in excess of $I_{1/2}$ (Supplementary Information). The growth of a community consisting of these strains is simulated by the specified consumer resource model (Supplementary Information). To simulate enrichment, all strains are started with identical biomass at fixed nitrate concentration and grown for $72\,\mathrm{h}$. We then simulate an 8-fold dilution into $2\,\mathrm{mM}$ nitrate. This repeated for a total of 12 cycles. (**B**) Simulation results are shown. Ecological outcomes for a parameter scan across the curvature parameter $k$ and toxicity parameter $I_{1/2}$ are classified by the peaks in the relative abundance distributions of the phenotypes after 12 simulated growth cycles. Increase of the parameter $I_{1/2}$ causes a decrease in toxicity, which corresponds to an increase in pH. Purple region (I) corresponds to coexistence of two strains capable of both nitrate and nitrite reduction ($r_A > 0$ and $r_I > 0$), while the orange region (II) corresponds to the dominance of a single strain. (**C**) Relative abundance distribution for the choice of parameters marked by the "x" in region I (B). Two generalists are enriched, one that does not die in the presence of high nitrite concentrations that grows preferentially on nitrite, and another that dies in the presence of high nitrite that grows preferentially on nitrate. This qualitatively reproduces the experimental enrichment outcome at pH 6.0. (**D**) Relative abundance distribution for the choice of parameters marked by the "x" in region II (B). One generalist is enriched, qualitatively reproducing the experimental enrichment outcome at pH 7.3. (**E**) Simulated co-culture enrichment of the two phenotypes drawn from the peaks of the relative abundance distribution shown in (C). The relative abundance of the strain with higher $r_A$, marked by a blue cell in (C) since it has a Nar$^+$ phenotype, is plotted. These phenotypes coexist, with the Nar$^+$ phenotype growing to higher relative abundance. This is qualitatively consistent with the experimental results shown in main text Fig. 5E. (**F**) Simulated co-culture enrichment of the phenotype at the peak of the relative abundance distribution shown in (D), along with the phenotype with the maximal $r_A$ for which $r_I > 0$. The second strain was chosen as a Nar$^+$ phenotype proxy, since the enriched generalist at low toxicity corresponds to the Nap$^+$ phenotype at pH 7.3. The relative abundance of the Nar$^+$ phenotype is plotted, and it approaches 0. Thus the phenotype marked by the orange cell in (D) dominates at low toxicity. This is qualitatively consistent with the experimental results shown in main text Fig. 5F.

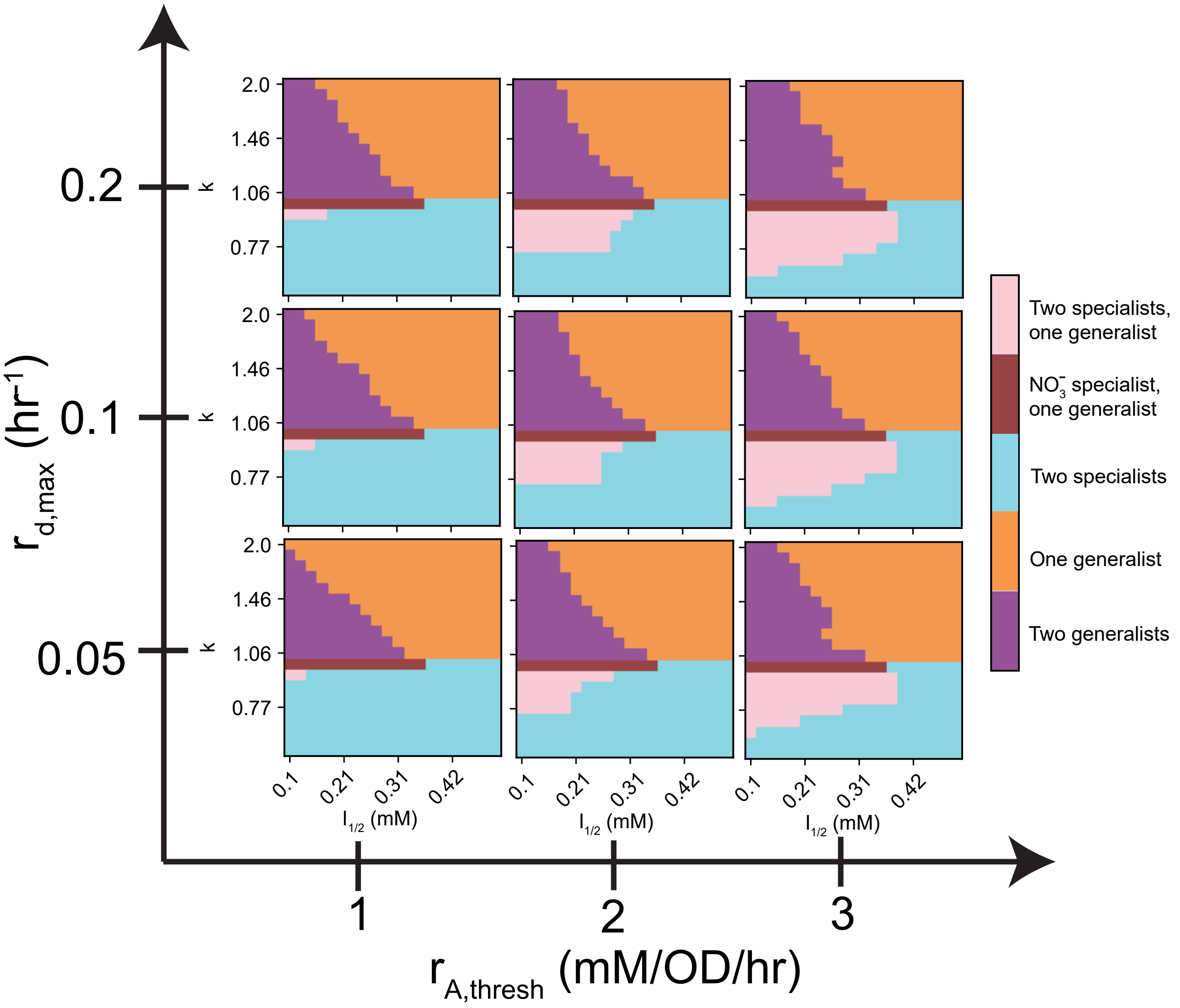


Figure S13: **Simulated enrichment outcomes dependence on parameters.** A scan of the simulation parameter space is shown. We varied the threshold $r_A$ where toxicity occurs ($r_{A,thresh}$), the threshold nitrite level for toxicity ($I_{1/2}$), and the maximum death rate ($r_{d,max}$). Colors indicate a classification of the endpoint community by the phenotype of the taxa at the peak of the relative abundance distributions as a function of the curvature parameter $k$ and toxicity parameter $I_{1/2}$. The results are not sensitive to variations in $r_{\mathrm{d,max}}$ and $r_{\mathrm{A,thresh}}$ around $0.1\,\mathrm{h}^{-1}$ and $2\,\mathrm{mM/OD/h}$, respectively, which were used in Fig. S12.

### Supplementary Tables

Table S1: Properties of soils used in enrichment experiments.

| Soil # | pH | Location | Latitude | Longitude | Date | Associated fig. |
|---|---|---|---|---|---|---|
| 1 | 7.5 | Meadowbrook Park, Urbana, IL | 40.081 | -88.203 | 11/1/20 | Fig. 2, Fig. S3 |
| 2 | 7.6 | Meadowbrook Park, Urbana, IL | 40.080 | -88.204 | 11/1/20 | Fig. 2, Fig. S3 |
| 3 | 7.2 | Meadowbrook Park, Urbana, IL | 40.080 | -88.205 | 11/1/20 | Fig. 2, Fig. S3 |
| 4 | 7.2 | Meadowbrook Park, Urbana, IL | 40.081 | -88.202 | 11/1/20 | Fig. 2, Fig. S3 |
| 5 | 7.3 | Meadowbrook Park, Urbana, IL | 40.081 | -88.202 | 11/1/20 | Fig. 2, Fig. S3 |
| 6 | 7.3 | Meadowbrook Park, Urbana, IL | 40.080 | -88.204 | 11/1/20 | Fig. 2, Fig. S3 |
| 7 | 6.9 | LaBagh Woods, Chicago, IL | 41.978 | -87.743 | 1/18/22 | Fig. S4 |
| 8 | 4.7 | Warren Woods, Three Oaks, MI | 41.837 | -86.631 | 2/21/22 | Fig. S4 |
| 9 | 5.9 | Warren Woods, Three Oaks, MI | 41.838 | -86.630 | 2/21/22 | Fig. S4 |
| 10 | 6.4 | Center Township, IN | 41.293 | -86.595 | 5/10/22 | Fig. S4 |
| 11 | 6.8 | Sauk Prairie State Recreation Area, Merrimac, WI | 43.362 | -89.735 | 4/23/22 | Fig. S4 |
| 12 | 5.8 | Barneveld Prairie State Natural Area, Barnevelde, WI | 42.987 | -89.899 | 4/23/22 | Fig. S4 |
| 13 | 7.5 | Bunker Hill, Niles, IL | 42.003 | -87.785 | 1/18/22 | Fig. S4 |

Table S2: Metagenome-assembled genome (MAG) taxonomy, statistics, and genotypes. Completeness and contamination estimated using CheckM (*111*). MAG annotation performed using RAST (*41*).

| pH | Soil sample | Taxon | Completeness (%) | Contamination (%) | narG | napA | nirS | nirK | norB | nosZ |
|---|---|---|---|---|---|---|---|---|---|---|
| 6 | 1 | Pseudomonadales | 99.86 | 0.216 | x | | x | | x | x |
| 6 | 2 | Pseudomonadales | 100 | 0.324 | x | | x | | x | x |
| 6 | 3 | Pseudomonadales | 99.83 | 0.126 | x | x | x | | x | x |
| 6 | 5 | Pseudomonadales | 100 | 0.566 | x | | x | | x | x |
| 6 | 6 | Pseudomonadales | 100 | 0.739 | x | x | x | | x | x |
| 7.3 | 1 | Pseudomonadales | 92.28 | 0.216 | x | | | | | |
| 7.3 | 3 | Pseudomonadales | 99.83 | 0.126 | x | x | x | | x | x |
| 7.3 | 4 | Pseudomonadales | 98.21 | 0.955 | x | | x | | x | x |
| 7.3 | 6 | Pseudomonadales | 100 | 0.216 | x | | x | | x | x |
| 6 | 1 | Rhizobiales | 98.66 | 1.66 | | x | | x | x | x |
| 6 | 2 | Rhizobiales | 99.61 | 0 | | x | | x | x | |
| 6 | 4 | Rhizobiales | 99.95 | 1.909 | | x | | x | x | x |
| 6 | 5 | Rhizobiales | 99.95 | 0.897 | | x | | x | x | |
| 6 | 6 | Rhizobiales | 99.61 | 1.273 | | x | | x | x | x |
| 7.3 | 1 | Rhizobiales | 89.18 | 2.344 | | x | | x | | x |
| 7.3 | 2 | Rhizobiales | 84.43 | 3.058 | | x | | x | x | x |
| 7.3 | 3 | Rhizobiales | 99.95 | 1.058 | | x | | x | x | x |
| 7.3 | 4 | Rhizobiales | 99.6 | 1.293 | | x | | x | x | x |
| 7.3 | 5 | Rhizobiales | 95.35 | 1.698 | | x | | x | x | x |
| 7.3 | 6 | Rhizobiales | 98.32 | 0.346 | | x | | x | x | x |
| 6 | 6 | Burkholderiales | 97.74 | 0.917 | x | x | | | | |
| 7.3 | 2 | Burkholderiales | 98.56 | 0.917 | x | x | | | | |
| 7.3 | 5 | Burkholderiales | 98.79 | 1.376 | x | | | | | |
| 6 | 3 | Enterobacteriales | 99.89 | 0.741 | x | | | | | |

| Study | Description | Results |
| --- | --- | --- |
| Lycus *et al.* (*27*) | 177 isolates from two soils, one neutral (pH 7.4 and one acidic (pH 3.7). 70 isolates were phenotyped and classified phylogenetically, 12 subjected to whole genome sequencing. Isolation is done by traditional plating methods. | 1. 22/23 isolates from acidic soil are Proteobacteria. Single additional strain is a Fermicute that performs DNRA.<br>2. 11/27 isolates from neutral soil are Proteobacteria. 13/16 remaining strains were DNRA or nitrate only reducers, primarily Actinobacteria.<br>3. All Actinobacteria are nitrate-only reducers.<br>4. 7/10 $NirK^+$ strains are from neutral pH, following Fig. 1H pattern. 3/4 $Nar^+$ and single $Nap^+$ strain from neutral pH, result is ambiguous. |

| | | |
|---|---|---|
| Graf, Jones, and Hallin. (*138*) | >600 genomes of denitrifiers (2012) with genotypes, phylogeny, and habitat annotated. Co-occurence of genotypes analyzed, no characterization of pH. Genomes taken from sequencing database. | 1. Majority of denitrifiers 369 / 652 are Proteobacteria.<br>2. from Lycus *et al.* Actinobacteria are the second most abundant phylum overall.<br>3. In soils NirK and NirS genes do not co-occur within the same genome, but they do (rarely) in other environments. |
| Gamble *et al.* (*28*) | 146 isolates from 23 soils and sediments sampled from across the globe. Standard isolation methods used, pH values range from 3.84 to 7.8. No genotype information are available. | 90% of the isolates are Proteobacteria, with the remaining 10% Flavobacteria. All soils except 1 (Table 3) yielded only Proteobacterial isolates. |
| Nishizawa *et al.* (*139*) | 38 isolates from three rice paddy soils ranging in pH from 5.6 to 6.8. Sophisticated single-cell isolation applied non-fermentable carbon sources, nitrate, and bacteriostatic antibiotics to microscopically identify and isolate active denitrifiers in soils (elongated cells) and isolate them individually. No genotype information available. | 33/38 isolates are Proteobacteria. |

Table S3: Meta-analysis of taxonomy of dominant denitrifiers in soils via culture-based methods.

Table S4: Properties of Cook Agronomy Farm soils used in topsoil incubation experiments (associated with Fig. 7).

| Soil_ID | pH | Latitude | Longitude | Sand (%) | Silt (%) | Clay (%) | Total carbon (%) | C:N ratio | $NO_3^-$ consumed (mM) |
|---|---|---|---|---|---|---|---|---|---|
| CE239 | 5.0 | 46.781 | -117.080 | 0.00 | 61.20 | 38.80 | 1.35 | 12.27 | 0.86 |
| CE201 | 5.3 | 46.781 | -117.086 | 0.00 | 58.70 | 41.30 | 1.57 | 13.07 | 1.28 |
| CE73 | 5.4 | 46.780 | -117.086 | 0.00 | 61.20 | 38.80 | 1.65 | 12.97 | 1.06 |
| CE277 | 5.8 | 46.782 | -117.084 | 0.00 | 61.20 | 38.80 | 1.19 | 12.65 | 0.84 |
| CE234 | 6.2 | 46.781 | -117.083 | 0.00 | 62.50 | 37.50 | 1.88 | 14.04 | 1.10 |
| CE229 | 6.3 | 46.781 | -117.085 | 0.00 | 58.80 | 41.20 | 1.20 | 12.73 | 0.86 |
| Neutral2 | 6.6 | 46.781 | -117.085 | 0.00 | 58.80 | 41.20 | 1.28 | 12.06 | 2.00 |
| Neutral5 | 6.8 | 46.781 | -117.085 | 0.00 | 63.70 | 36.30 | 1.31 | 13.79 | 2.00 |
| Neutral6 | 6.9 | 46.782 | -117.085 | 0.00 | 61.20 | 38.80 | 1.07 | 12.30 | 2.00 |
| Neutral3 | 7.1 | 46.781 | -117.085 | 0.00 | 60.00 | 40.00 | 1.09 | 12.56 | 2.00 |
